# Bispecific antibody targeting of lipid nanoparticles

**DOI:** 10.1101/2024.12.20.629467

**Authors:** Angelo Amabile, Matthew Phelan, Zhixin Yu, Pedro Silva, Adam Marks, Judit Morla-Folch, Moah Sohn, Gurkan Mollaoglu, Chiara Falcomata, Abraham J P Teunissen, Joshua D Brody, Yizhou Dong, Brian D. Brown

**Author notes:** These two authors contributed equally to this work (Matthew Phelan, Zhixin Yu).

## Abstract

Lipid nanoparticles (LNP) are the most clinically advanced non-viral gene delivery system. While progress has been made for enhancing delivery, cell specific targeting remains a challenge. Targeting moieties such as antibodies can be chemically-conjugated to LNPs however, this approach is complex and has challenges for scaling up. Here, we developed an approach to generate antibody-conjugated LNPs that utilizes a bispecific antibody (bsAb) as the targeting bridge. As a docking site for the bsAb, we generated LNPs with a short epitope, derived from hemagglutinin antigen (HA), embedded in the PEG component of the particle (LNP^HA^). We generated bsAb in which one domain binds HA and the other binds different cell surface proteins, including PD-L1, CD4, CD5, and SunTag. Non-chemical conjugation of the bsAb and LNP resulted in a major increase in the efficiency and specificity of transfecting cells expressing the cognate target. LNP/bsAb mediated a 4-fold increase in *in vivo* transfection of PD-L1 expressing cancer cells, and a 26-fold increase in ex vivo transfection of quiescent primary human T cells. Additionally, we created a universal bsAb recognizing HA and anti-rat IgG2, enabling LNP tethering to off-the-shelf antibodies such as CD4, CD8, CD20, CD45, and CD3. By utilizing a molecular dock and bsAb technology, these studies demonstrate a simple and effective strategy to generate antibody-conjugated LNPs, enabling precise and efficient mRNA delivery.

## Introduction

Lipid nanoparticles (LNP) have transformed the field of RNA therapeutics by providing an efficient means for *in vivo* RNA delivery^1^. The first FDA approved siRNA drug and BioNTech and Moderna’s COVID-19 mRNA vaccines utilize LNP technology for RNA delivery. LNP are lipid-based vesicles with a diameter ranging from 50 to 200 nm^2^. They typically consist of ionizable lipids, cholesterol, phospholipids, and polyethylene glycol (PEG)-lipids^1^. At acidic pH, the tertiary amine of the ionizable lipid interacts with the negatively charged RNA, facilitating its encapsulation in the LNP. Cholesterol provides structural integrity and stability to the vesicle, while PEG-lipids and phospholipids form the surface layer. PEG also creates the hydrated layer around the LNP, preventing aggregation and ensuring colloidal stability *in vitro* and *in vivo*^3^.

Upon systemic administration, PEG-lipids progressively shed from the LNP surface, allowing serum proteins to absorb onto the particles^4,5^. This process imparts tissue-directed targeting capabilities to the LNPs as the serum proteins influence the tissue tropism of the LNP^6,7^. Notably, many LNPs become bound by apolipoprotein E (APOE) and target the liver via the low-density lipoprotein receptor (LDL-R), making LNPs an attractive vehicle for the treatment of hepatic diseases^8^. However, many RNA therapeutics require delivery of the RNA to specific cell types. For example, to eliminate tumors, an RNA encoding a toxic payload must be targeted solely to cancer cells and avoid healthy cells and tissues. Targeted LNPs can also extend the cell and tissue tropism of RNA delivery; for instance, by increasing the efficiency of delivery to otherwise poorly transfected cell types, such as B cells or hematopoietic stem cells (HSC)^7,9^, and this can have utility not only in vivo, but also in vitro for engineering cells prior to transplant.

Extensive efforts have been made to generate targeted LNP therapeutics to broaden cell types transfected and to improve therapeutic index^7^. Different strategies have been developed for the generation of tissue/cell specific LNP. Modifying the ratio of the different lipids and inclusion of accessory lipids can have a profound impact on tissue tropism. An example is the Selective Organ Targeting (SORT) system^10^, which includes a lipid such as DOTAP, to vary the net charge of the LNP, and in doing so, increase the tropism of the LNP to extrahepatic organs such as the spleen and lung. Another approach involves engineering the LNP surface with a ligand that binds specific cell surface molecules. This can be achieved by substituting PEGylated lipids with PEG molecules linked to targeting moieties such as peptides^11–13^, antibodies^14^, aptamers^15^ or small molecules^16^. As example, peptide guided LNPs have been used to deliver mRNA to the neural retina of rodents and non-human primates^11^. Typically, PEGylated distearyl lipids (C18) are used instead of dimyristoyl lipids (C14) for these LNPs since the desorption rate of distearyl lipids in the serum is lower than dimyristoyl lipids which ensures that targeting moieties can be present on the surface of LNPs to the utmost extent upon injection^4^.

Antibody-targeted LNPs (Ab-LNP) provide a rationale and potent means for enhancing delivery to specific cell types. For example, LNPs have been conjugated to anti-CD4 or anti-CD5 to target delivery to T cells^17,18^, anti-CD38 to target multiple myeloma cells^19^, anti-PECAM1 and anti-CD31 to target endothelial cells^14^, and anti-CD117 to target hematopoietic stem cells (HSC)^20,21^. Since the antibodies are not well tolerated to organic solvents used in the LNP preparation, they are chemically conjugated to the LNP after RNA encapsulation. The most widely used antibody conjugation method is the thiol-Maleimide reaction^14,17,22^. Maleimide functionalized LNPs are first prepared using the standard method by incorporating Mal-PEG-DSPE in lipid mixtures. Then, antibodies are subjected to reducing agents to expose free reactive thiol groups for the conjugation step. For antibodies whose thiols are unavailable or absent, a protein modification agent, N-succinimidyl S-acetylthioacetate (SATA), is normally utilized to introduce external thiol groups into protein molecules. Despite great achievements, the nature of the chemical conjugation method makes Ab-LNP difficult to standardize and manufacture at higher scale^23–25^. High batch-to-batch variability, potential affinity reduction of antibodies, and the requirement for individual optimization for each antibody reagents have so far limited the approach to preclinical experimental applications^23^. Though there are ongoing efforts to bring the approach to clinical applications, even for ex vivo cell manipulation, the technology has still remained niche.

Bispecific antibodies (bsAb) have an increasing number of applications, including in the clinic, such as in cancer immunotherapy and for treatment of hemophilia^26,27^. Advances in engineering technologies have significantly streamlined their manufacturing, reducing production complexity and costs while enabling scalable and reproducible generation of these versatile molecules^28^. Their smaller size and lack of an antibody constant region enhance their flexibility, enabling better tissue penetration, reduced immunogenicity, and improved adaptability for innovative targeting strategies.

We reasoned that bsAb could be leveraged to redirect LNPs toward specific cell populations without the need for chemical conjugation. To achieve this, we developed a method to couple bsAb to LNPs by incorporating a short linear epitope into the PEG component of the LNP. One arm of the bsAb recognizes this epitope, while the other arm is engineered to target a specific surface membrane protein expressed on the cells of interest. Assembly of the bsAb/LNP complexes occurs in physiological conditions without the need of coupling reagents and purification, and results in uniform particle generation. We show that bsAb/LNP increase cell specific targeting both at saturating and limiting doses of LNP while reducing off-target activity both *in vitro* and *in vivo.* LNP/bsAb complexes also increase delivery efficiency to otherwise difficult-to-transfect cell types such as quiescent human T cells. Overall, we generated a valid alternative to traditional Ab-LNP conjugation that can be easily implemented for the generation of cell-targeted LNPs.

## Results

### Generation of linear epitope binding bispecific antibody as a tool compound

BsAb have many applications^27^, but there is no available bsAb tool compound that can recognize model epitopes and could be used for rapid testing of bsAb functions, including for LNP conjugation. To meet this need, we generated a bsAb recognizing linear epitopes from HA and SunTag. This was done by cloning two single chain variable fragments (scFv) derived from the antigen binding domain (Fab) sequences of SunTag and HA antibodies, separated by the 218s linker^29^ (bsAb^HA-SunTag^) (Fig. 1a). The HA^30^ and SunTag^31^ were chosen because they are short linear epitopes that can easily be fused to other molecules, including potentially PEG, and the scFV have been engineered to increase solubility. We transfected Expi293 cells with our expression plasmid and allowed 72 hours for sufficient production of our construct. Supernatant was collected and concentrated by nickel column purification and protein integrity was verified by Western-Blot (Fig. S1a and S1b).

**Figure 1.**
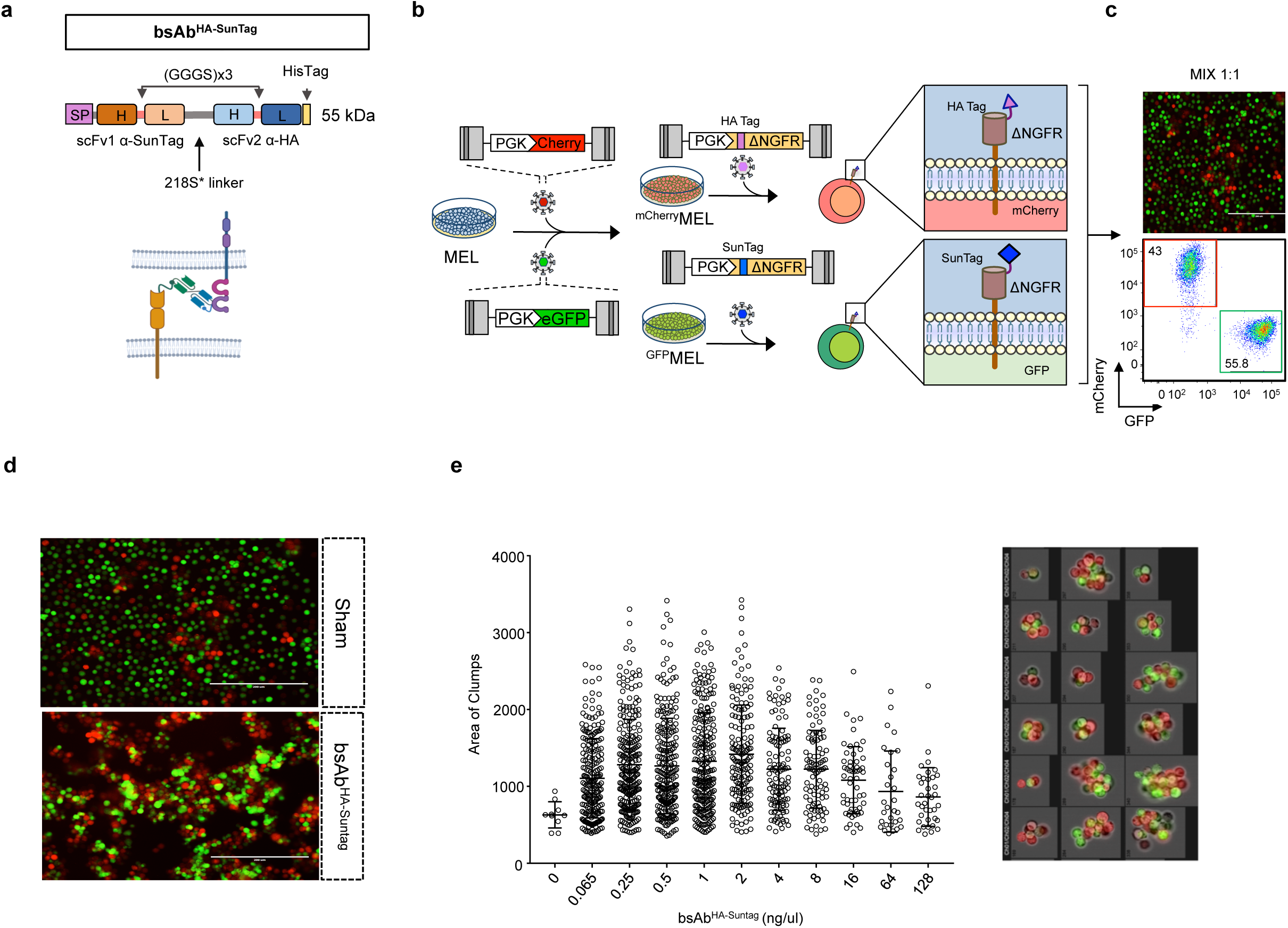
Generation of a linear epitope binding bispecific antibody. a. Schematic of bsAb^HA-SunTag^ and depiction of activity in bringing epitope expressing cell types together. SP:signal peptide, scFV: single chain variable fragment. b. Schematic outlining generation of the cell populations expressing the targets of the bsAb^HA- SunTag^ used in the experimental conditions. MEL leukemia cells were transduced with a lentiviral vector (LV) expressing mCherry or GFP. MEL^mCherry^ and MEL^GFP^ were subsequently transduced with a LV expressing delta-NGFR^HA^ and delta-NGFR^Suntag^ respectively. MEL^mCherry/NGFR-HA^ and MEL^GFP/NGFR-Suntag^ were mixed in 1:1 ratio. c. The relative frequency of MEL^mCherry/NGFR-HA^ and MEL^GFP/NGFR-Suntag^ cells was assessed by florescent microscopy and flow cytometry. Shown is a representative image (20x) of the cell mix (top) and dot plot from the flow cyometry (bottom). d. Microscopy image of MEL^GFP/NGFR-Suntag^ and MEL^mCherry/NGFR-HA^ cell cultures incubated with 1 mg/ul of purified bsAb^HA-SunTag^ or unrelated bsAb. Image is representative from 3 independent cell-cell aggregation experiments. e. Quantification (left histogram) and visualization (representative images on the right) of clumps of MEL^eGFP/NGFR-Suntag^ and MEL^mCherry/NGFR-HA^ cells measured by Amnis® Image-Stream analysis. Experiment was repeated 3 independent times.

To test the functionality of both binding arms of the bispecific, we developed a cell-cell aggregation assay to assess the ability of the bsAb^HA-SunTag^ to bring HA and SunTag expressing cells together, similar to how a T cell engager bsAb functions. We engineered two lines of Mouse Erythroleukemia (MEL) cells; one expressing enhanced green florescent protein (GFP) and a truncated neurotrophic growth factor receptor (ΔNGFR) with SunTag fused to the N-terminus (MEL^GFP,SunTag^) and the other MEL line expressing mCherry florescent protein (mCherry) and ΔNGFR with HA fused to the N-terminus (MEL^Cherry,HA^) (Fig. 2b). We co-cultured 10^5^ cells of each line in 1:1 ratio and added our bsAb^HA-SunTag^ from a dose range of 0.065 – 128 ng/ul. Cells were then incubated for 48 hours, after which they were analyzed by fluorescent microscopy and Amnis® ImageStream MkII flow cytometry.

**Figure 2.**
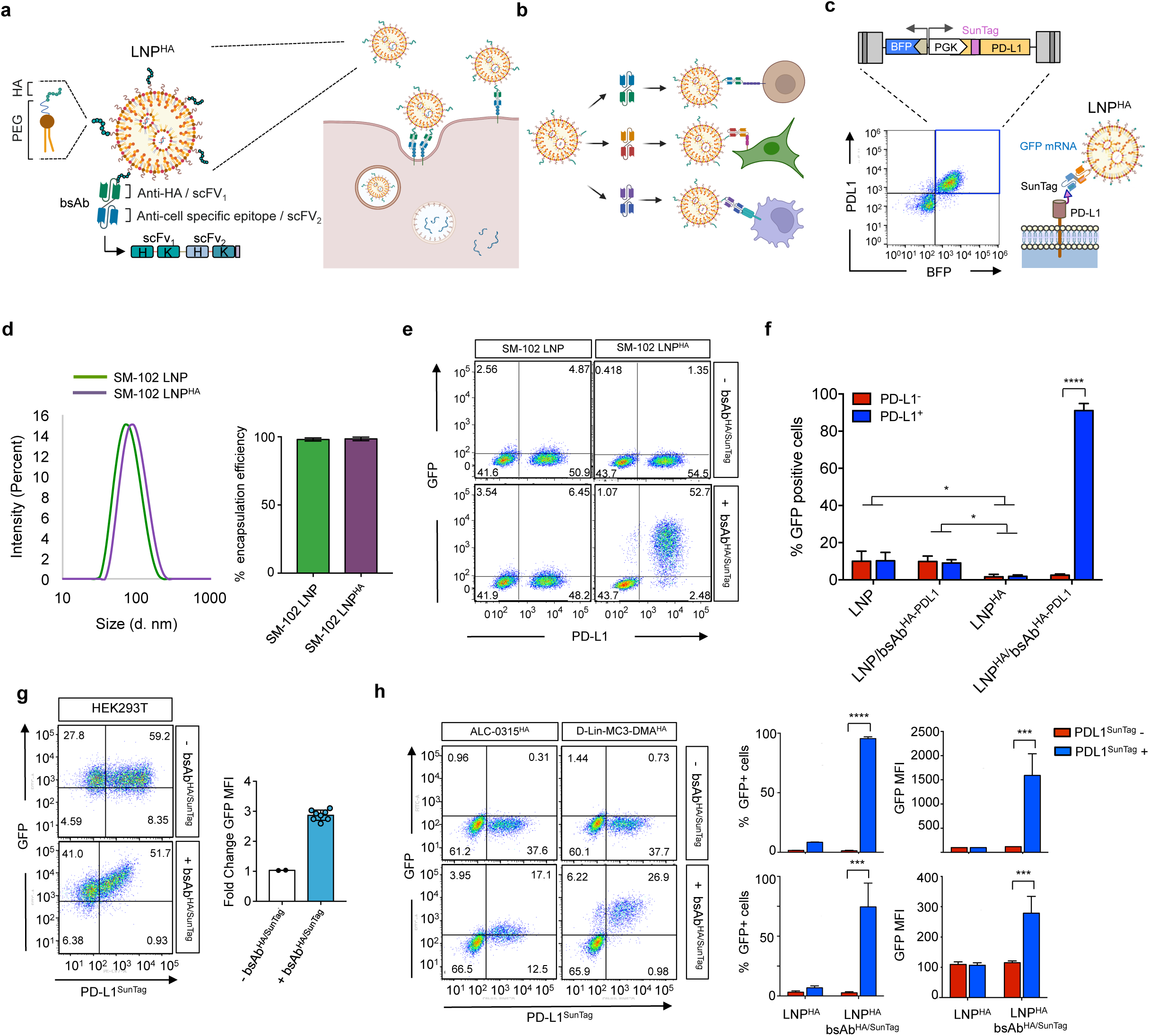
bsAb-coupled LNP mediates enhanced ligand-specific cell transfection. a. Schematic of the LNP^HA^ with bound bsAb (left) and depiction of bsAb-mediated uptake through ligand binding. b. Schematic showing how LNP^HA^ can be redirected to different cell types with different bsAb. c. Schematic showing the bidirectional lentiviral vector (Bid.LV) used to generate PD-L1^Suntag^/BFP reporter cell line (top). Representative dot-plot of human erythroleukemia cells (K562) transduced with Bid.LV PD-L1^Suntag^/BFP, K562^PD-L1/SunTag^ (Bottom). d. Size analysis by DLS of SM-102 LNP (green) and SM-102 LNP^HA^ (purple) particles (left). Histograms showing mRNA encapsulation efficiency in LNP and LNP^HA^ as measured by RiboGreen assay (right). Graphs are representative of all LNP-mRNA particles generated in the study. e. Flow cytometry analysis of GFP and and PD-L1 expression in K562^PD-L1/SunTag^ cells transfected with 10 ng of GFP mRNA in LNP or LNP^HA^ with or without bsAb^HA-Suntag^. Shown are representative dotplots. Data representative of 5 independent experiments, with n=3 biological replicates for each experiment. f. Graphs show the mean±s.d. percentage of GFP-positive K562^PD-L1/SunTag^ cells from (e). Analysis of GFP expression was performed 24 hours post-transfection. Two-way anova and Tukey’s multiple comparison post-test, **p<0.001; ****p<0.00001 (n=3). Data representative of 5 independent experiments. g. Flow cytometry analysis of GFP and and PD-L1 expression in 293T^PD-L1/SunTag^ cells transfected with 10 ng of GFP mRNA in LNP or LNP^HA^ with or without bsAb^HA-Suntag^. Representative dotplots shown (n=3-8, 3 independent experiments) (left). Graph (right) shows fold change of GFP mean florescence intensity (MFI) of PD-L1^Suntag^ positive cells treated with bsAb^HA-Suntag^ compared to untreated cells. h. Flow cytometry analysis of GFP and and PD-L1 expression in K562^PD-L1/SunTag^ cells transfected with 10 ng of GFP mRNA in Dlin-MC3 LNP^HA^ or ALC-0315 LNP^HA^ in absence (left column) or presence (right column) of bsAb^HA-Suntag^. Analysis performed 24 hours post transfection. i. Graphs showing percentage (left) and MFI (right) of GFP positive cells from (h). Statistical analysis: two-way anova and Tukey’s multiple comparison post-test,; ***p<0.0001 (n=3) Data representative of one independent experiment.

Upon visual inspection we could see in sham treated conditions that GFP+ and mCherry+ cells were mostly separated single cells, whereas in cultures with the bsAb^HA-SunTag^ there was a noticeable degree of mCherry+ and GFP+ cell-cell aggregation (Fig. 1c). We confirmed a significant increase in cell aggregation in the bsAb^HA-SunTag^ cultures using ImageStream analysis, by quantifying the total area of GFP and mCherry double positive clusters (Fig. 1d). Our bsAb^HA-SunTag^ appeared to work best within a range from 0.065 ng/ul to 1.875 ng/ul, with a predictable reduction in efficiency at higher doses (>3.75 ng/ul), indicative of a Hook effect in which all antigen is bound by either antigen binding domain of an antibody, preventing cells from coming into contact with one another.

To test how other linear epitopes might perform as a bsAb target, we substituted the scFv of SunTag with the scFv of a Flag Tag antibody (bsAb^HA-Flag^). Testing in our cell aggregation assay indicated that bsAb^HA-Flag^ also mediated an increase in aggregation of target expressing cells, in this case MEL^GFP,Flag^ and MEL^Cherry,HA^ cells (Fig. S1c,d). These results demonstrate the generation and validation of linear epitope targeted bsAb bind their targets and can promote cell-cell clustering. They also establish new tool compounds for bsAb studies which recognize commonly used fusion tags.

### BsAb mediate cell-specific targeting of LNPs

We sought to test if we could attach the bsAb^HA-SunTag^ to the surface of an LNP modified to contain the HA peptide, and use these particles to enhance the transfection SunTag expressing cell (Fig. 2a, 2b). We engineered HEK293T cells and K562 leukemia cells to co-express a blue fluorescent protein (BFP) reporter and a recombinant fusion of human PD-L1 with the SunTag in the N-terminal (PD-L1^SunTag^) (Fig. 2c). We chose PD-L1 because it is a molecule that is highly upregulated on cells within tumors, and serves to promote T cell exhaustion and immune evasion^32^. We used BFP as a reporter to enable us to monitor LNP^HA^ transfection of PD-L1^SunTag^ positive cells without the need for staining for PD-L1, whose signal can be impaired by endocytosis of the receptor and/or steric hindrance of the bound bsAb^HA-SunTag^. The cells were confirmed to co-express BFP and PD-L1 (Fig. 2c).

We generated in vitro transcribed (IVT) modified mRNA^33^ encoding for GFP and encapsulated in an SM-102-based LNP^34^ in which 15% of total PEG was substituted with PEG^HA^, a PEG lipid with the short HA peptide covalently attached. We then incubated the LNP^HA^ with bsAb^HA-SunTag^. The resulting particles were only slightly bigger in size compared to standard LNP (84.3±8.8 and 73.80±0.628 respectively) and showed similar PDI (0.11± 0.03 and 0.15±0.05) and mRNA encapsulation efficiency (Fig. 2d, S2a).

We mixed K562^PDL1-SunTag^ and K562 expressing cells in a 1:1 ratio to have target+ and target-cells in the same culture. Transfection with LNP or LNP^HA^ led to little to no GFP expression in the cells, consistent with the fact that at the utilized LNP-RNA dose, K562 cells are poorly transfected by LNPs. (Fig. 2e, 2f). Instead, when the cells were transfected with LNP^HA^ that was pre-incubated with bsAb^HA-SunTag^, >90% of PD-L1+/SunTag+ cells became GFP+, and at much higher GFP levels than control-treated cells (Fig. 2e, 2f, S2b). Transfection was remarkably specific as only ∼1% of PD-L1-negative/SunTag-negative cells became GFP+ in the same culture. Target cell transfection was dependent on bsAb being bound to the LNP^HA^ particle, since the K562^PDL1-SunTag^ were not transfected by SM-102 LNP (no PEG^HA^) that had been pre-incubated with bsAb^HA-SunTag^.

As we found that the bsAb-coupled LNP not only increased specificity but also delivered more RNA per cell, as indicated by the higher GFP MFI in transfected cells (Fig. 2e), we wanted to determine if the bsAb/LNP would enhance transfection of cells already permissive to LNP-RNA delivery. To test this, we transfected 293T^PDL1-SunTag^ cells, and once again we used a 1:1 mix of target expressing and non-expressing cells (i.e. 293T: 293T^PDL1-SunTag^). With LNP^HA^, ∼90% of cells were transfected and expressed GFP at an MFI of 2,100. When the cells were transfected with LNP^HA^/bsAb^HA-SunTag^, a similar percent of cells were transfected, but the cells expressing PD-L1^SunTag^ had a GFP MFI of 5,880, indicating that even in permissive cells, bsAb-mediated targeted transfection increased RNA delivery by ∼3-fold (Fig. 2g). This data indicates that our bsAb/LNP-targeting technology allows transfection of otherwise refractory cell lines and increases the level of mRNA expression at saturating doses in transfection-permissive cells.

To determine if the efficacy of LNP^HA^ targeting was independent from LNP chemical composition, we generated LNP^HA^ using different sets of lipids, based on different FDA approved LNP formulations. Specifically, we encapsulate GFP-encoding mRNA in LNP^HA^ containing Dlin-MC3-DMA (Alnylam’s Onpattro-like) and ALC-0315 (Pfizer mRNA vaccine-like). When K562^PDL1-SunTag^ were transfected with either of the additional LNP^HA^ formulations coupled with bsAb^HA-SunTag^, we once again found that there was efficient and specific delivery to target-expressing cells, as indicated by GFP-positivity in cells expressing PD-L1^SunTag^ (Fig. 2h). This data indicates that bsAb coupling can be used to enhance the targeting of different LNPs formulations.

To determine the optimal bsAb amount to couple with the LNP, we incubated 10 ng of LNP^HA^ with a concentration of bsAb^HA/SunTag^ ranging from 1 ng up to 200 ng, and then transfected K562^PDL1-SunTag^ cells. We observed maximum activity, measured as MFI of PD-L1+ cells, with 40 ng of bsAb^HA/SunTag^, with reduced transfection efficiency at lower and higher bsAb doses (Fig. S2c). At 1:1 weight ratio and lower bsAb:LNP^HA^ ratios, there was a modest increase in LNP^HA^ size (∼10 nm) indicating binding of bsAb to the HA tag on the surface of the LNP^HA^. At highest bsAb concentration (100 and 40 ng) there was a 3-fold and 2-fold increase in LNP^HA^ size, respectively, most likely reflecting aggregation of LNP^HA^ due to an excess of bsAb (Fig. S2d). We also titrated the range of PEG^HA^ that can be incorporated in the LNP preparation, evaluating a gradient in which PEG^HA^ accounts from 7.5% to 30% of total PEG. The higher the PEG^HA^ content, the bigger the size and PDI values of the LNP^HA^ (Fig. S2e) with a concomitant decrease in mRNA encapsulation efficiency and recovery rate (data not shown). We selected 15% PEG^HA^ substitution rate and 1:1 bsAb:LNP weight ratio for all future experiments, since at this dose particles retained PDI values and mRNA encapsulation efficiency similar to standard LNP and showed high transfection efficiency with a modest size increase (∼10 nm) (Fig. S2f).

### LNP^HA^/bsAb complexes increase targeting of PD-L1 positive cancer cells

To assess the efficiency of LNP^HA^/bsAb targeting to a therapeutically relevant molecule, we generated a new bsAb in which we substituted the anti-SunTag scFv with the scFV of Atezolizumab, a monoclonal antibody that recognizes both human and murine PD-L1, to create bsAb^HA-PDL1^. BsAb^HA-SunTag^ and bsAb^HA-PDL1^ bind to different epitopes of PD-L1^SunTag^ (Fig. 3a).

**Figure 3.**
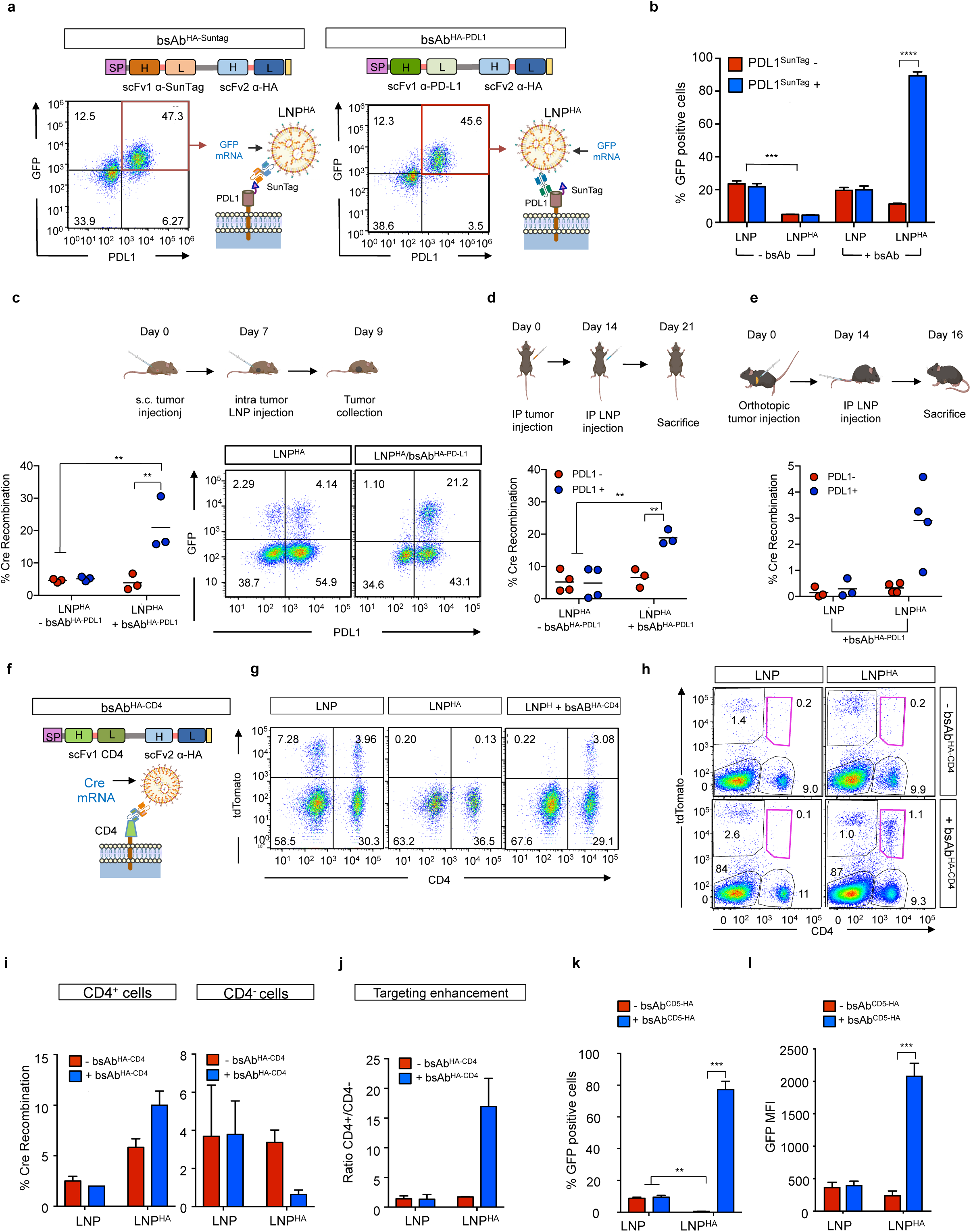
LNP^HA^/bsAb increases in vivo targeting of PD-L1+ tumor and CD4 T cells. a. Schematic of the bsAb^HA-Suntag^ and its binding domain on K562^PD-L1-SunTag^ (bottom left) and bsAb^HA/PD-L1^ (top right) and its binding domain on K562^PD-L1-SunTag^ (bottom right). Flow cytometry analysis of K562^PD-L1-SunTag^ treated with 10 ng of GFP mRNA LNP^HA^ incubated with either bsAb^HA-Suntag^ or bsAb^HA-PD-L1^. GFP and PD-L1 levels were measured after 24 hours. b. Graphs show the mean±s.d. percentage of GFP+ K562^mPDL1^ transfected with 20 ng GFP mRNA LNP or LNP^HA^ +/-bsAb^HA/PD-L1^ (n=3 biological replicates, 2 independent experiments). Two-way anova and Tukey’s multiple comparison post-test, ***p<0.0001; ****p<0.00001. c. Graphs show the percentage of transfected PD-L1+ and PD-L1- B16F10^dsRed-LSL-GFP^ melanoma cells following intratumoral injection of melanomas with matching concentrations of Cre mRNA encapsulated in LNP^HA^ or LNP^HA^+bsAb^HA-PD-L1^. Analysis was performed by flow cytometry. Each dot is a separate mouse. Two-way anova and Tukey’s multiple comparison post-test, **p<0.001. Schematic of the experiment shown (top). d. Graphs show the percentage of transfected PD-L1+ and PD-L1- ID8^dsRed-LSL-GFP^ ovarian cancer cells following i.p. injection of ovarian tumor bearing mice with matching concentrations of Cre mRNA encapsulated in LNP^HA^ or LNP^HA^+bsAb^HA-PD-L1^. Analysis was performed by flow cytometry. Each dot is a separate mouse. Two-way anova and Tukey’s multiple comparison post-test, **p<0.001. Schematic of the experiment shown (top). e. Graphs show the percentage of transfected PD-L1+ and PD-L1-KC^dsRed-LSL-GFP^ pancreatic cancer cells following i.p. injection of pancreatic tumor bearing mice with matching concentrations of Cre mRNA encapsulated in LNP^HA^ or LNP^HA^+bsAb^HA-PD-L1^. Each dot is a separate mouse. Two-way anova and Tukey’s multiple comparison post-test, **p<0.001. Schematic of the bsAb^HA-CD4^ (top left) bound to its target, CD4, on murine T cell. f. Schematic of the bispecific antibody recognizing HA and CD4. g. Flow cytometry analysis of tdTomato and CD4 expression on total splenocytes isolated from Ai14 mice and transfected with 50 ng Cre mRNA encapsulated in LNP, LNP^HA^, LNP^HA^+bsAb^HA-CD4^. Cultures included IL-2 and anti-CD3/CD28 beads to activate/expand T cells. Shown are representative dotplots. h. Flow cytometry analysis of splenocytes isolated from Ai14 mice i.v. injected with Cre mRNA encapsulated in LNP or LNP^HA^ and coupled with or without bsAb^HA-CD4^. Spleens collected 3 days post injection. Dotplots are representative of n=3 mice group, 2 independent experiments. i. Graphs show percent transfection of CD4+ and CD4-splenocytes from (h) (n=3 mice, 2 independent experiments). j. Graphs show in vivo targeting efficiency of the different LNP formulations calculated by the ratio of tdTomato+ CD4+ to CD4- splenocytes from (h). k. Graphs show the percentage of GFP+ resting human T cells treated with 200 ng GFP mRNA encapsulated in LNP or LNP^HA^ with or without bsAb^HA-CD5^. Expression was measured 24 hours post transfection. Two-way anova and Tukey’s multiple comparison post-test, **p<0.001; ***p<0.0001(n=4). Data representative of 3 independent experiments. l. Graphs showing the GFP MFI of resting human T cells treated as in (k). Two-way anova and Tukey’s multiple comparison post-test, ***p<0.0001(n=4). Data representative of 3 independent experiments.

We encapsulated GFP mRNA in SM-102 LNP and LNP^HA^ and incubated with or without bsAb^HA-SunTag^ and bsAb^HA-PDL1^. As before, LNP^HA^/bsAb^HA-SunTag^ mediated specific and efficient delivery to SunTag expressing K562^PDL1-SunTag^ cells (Fig. 3a). Similarly, bsAb^HA-PDL1^ promoted cell-specific transfection, with ∼95% of PD-L1+ cells becoming GFP+ when treated with LNP^HA^/BsAb^HA-PDL1^ and only low levels of off-target transfection of PD-L1-negative cells (Fig. 3a). There was a 1.4-fold increase in GFP MFI of cells treated with LNP^HA^/bsAb^HA-PDL1^ compared to LNP^HA^/bsAb^HA-SunTag^ indicating that targeting a natural ligand, PD-L1, did not negatively impact targeting activity and even improved transfection efficiency of LNP^HA^ (Fig. S3a).

To verify the ability of bsAb^HA-PDL1^ to also recognize mouse PD-L1, we generated K562 expressing murine PD-L1 (K562^mPDL1^). When LNP^HA^ and bsAb^HA-PDL1^ were coupled and added to cultures of K562^mPDL1^ in which ∼45% of cells expressed mPD-L1 on their surface, there was a major increase in LNP transfection of the mPDL1+ cells (Fig. 3b). As in the previous experiments conducted on K562^PDL1-SunTag^, LNP^HA^ particle alone were less effective than standard SM-102 LNP in transfecting K562^mPDL1^ (∼5% vs ∼20% respectively) (Fig. 3b). However, when LNP^HA^ was coupled with BsAb^HA-PDL1^, >90% of mPD-L1+ cells were transfected, and there was a 3-fold increase in GFP expression levels compared to SM-102 LNPs (Fig. 3b, S3b).

GFP mRNA provides cellular resolution of transfection efficiency but the instability of the mRNA results in GFP expression decreasing over time. As a more sensitive system for assessing transfection efficiency, we utilized a Cre-loxP recombination system. Using a lentiviral vector, we engineered ID8 ovarian cancer cells to encode a dsRed-STOP-loxP-GFP expression cassette. This results in the cells stably expressing dsRed florescent protein, but when cells express Cre recombinase the dsRed is excised and GFP is expressed (Fig. S3c).

We encapsulated Cre encoding mRNA into SM-102 LNP or SM-102 LNP^HA^ +/- bsAb ^HA-PDL1^. In vitro transfection of the ID8 cells with SM-102 LNP (+/-bsAb), resulted in 25% of the PD-L1+ cells becoming dsRed negative. In contrast, cells treated with SM-102 LNP^HA^/bsAb^HA-PDL1^ were ∼97% dsRed negative, indicating the LNP^HA^/bsAb complexes efficiently directed the particles to the mPD-L1 expressing cells (Fig. S3d).

### LNP^HA^/bsAb enhances *in vivo* transfection of targeted cells

Next, we sought to assess efficiency of targeted delivery *in vivo*. For these experiments we utilized the dsRed-STOP-loxP-GFP ID8 ovarian cancer cells, and also engineered KPC pancreatic cancer cells^35^ and B16F10 melanoma cells with the same reporter construct. The cells were engineered to express blue fluorescent protein (BFP) so they could be tracked *in vivo* and mPD-L1 (Fig. S3e).

To assess targeted transfection in a model of melanoma, we injected the reporter B16F10 melanoma cells (BFP+dsRed+GFP-mPD-L1+) subcutaneously (s.c.) into wildtype mice. We encapsulated Cre mRNA into the LNP^HA^ particle and incubated +/- bsAb^HA-PDL1^, and when tumor size reached 5 mm in diameter, we injected the LNPs into the tumor. After 3 days, we harvested the tumors and performed flow cytometry to assess the percent of BFP+, GFP+, and mPDL1+ cells. In mice that received LNP^HA^ with no bsAb, <4% of cells were GFP+. In contrast, in tumors injected with LNP^HA^/bsAb^HA-PDL1^, >20% of mPDL1+ cancer cells were GFP+ (Fig. 3c).

As a more challenging delivery context, we assessed bsAb-mediated LNP targeting to disseminated tumors. To do this, we used the ID8 ovarian cancer model. The dsRed+ BFP+ mPD-L1+ ID8 cancer cells were mixed with the parent ID8 cancer cells (no BFP, no dsRed, no mPD-L1), in a 1:1 ratio, and then intraperitoneal (i.p.) injected into immunocompetent mice. This results in disseminated tumors throughout the peritoneal cavity and models metastatic ovarian cancer^36^. After 10 days, when tumors had engrafted and formed macroscopic lesions, we injected LNP^HA^ +/- bsAb^HA-PDL1^ carrying Cre mRNA into the peritoneal cavity. Once the mice developed ascites (indicative of large tumors), we sacrificed the animals and analyzed the BFP+ cancer cells for expression of GFP by flow-cytometry (Fig. 3d). Without the bsAb, the LNP^HA^ mediated ∼5% transfection of the cancer cells and this was similar between PD-L1- and PD-L1+ cancer cells. Impressively, with LNP^HA^/bsAb^HA-PDL1^, ∼20% of PD-L1+ cancer cells were transfected, whereas only ∼6% of PD-L1-negative cancer cells were transfected in the same animals, indicating that the bsAb coupled LNP mediated a 3-fold increase in transfection efficiency specifically to target expressing cells *in vivo* (Fig. 3d).

Lastly, we tested targeted delivery to pancreatic tumors, using an orthotopic model of the disease. We injected KC pancreatic cancer cells^37^, which carry an activating Kras^G12D^ mutation, into the pancreas of mice and allowed tumors to grow (Fig. 3e). After 2 weeks, when tumors were significant size, we injected SM-102 LNP or LNP^HA^/bsAb^HA-PDL1^ encapsulating Cre mRNA into the peritoneal cavity, which leads to vascular dissemination of the particle. Two days later we harvested the tumors and analyzed BFP and GFP expression by flow cytometry. Though overall transfection efficiency was modest, unsurprisingly since these tumors grow in the pancreas and are very poorly vascularized, the bsAb coupled LNP was able to mediate a 3-fold increase in cancer cell delivery. Importantly, transfection was specific for PD-L1+ cancer cells (Fig. 3e), with as many as 5% of PD-L1+ pancreatic cancer cells transfected from systemic delivery of the LNP.

Altogether, these data demonstrate that the binding of the bsAb to LNP^HA^ is preserved *in vivo*, and that this non-chemical conjugation approach is able to target the LNP to cancer cells expressing a specific molecule, in this case PD-L1, and enhance uptake and expression of a locally or systemically delivered mRNA to cells *in vivo*.

### BsAb-coupled LNPs enhance efficiency and specificity of RNA delivery to primary mouse and human T cells

T cells have emerged as an important therapeutic target of LNP-RNA^22,38^. Several groups have reported successful *in vivo* targeted delivery to T cells using antibody-conjugated LNPs^17,18^. We set out to determine if bsAb coupled LNPs could also mediate delivery to T cells. To test this, we generated a bsAb comprised of the HA-targeting scFv and an scFv specific for murine CD4 (bsAb^HA-CD4^, Fig. 3f). We encapsulated Cre mRNA into LNP^HA^ and incubated with bsAb^HA-CD4^ to decorate the LNP particle. Total splenocytes were isolated from Ai14 mice, which encode a STOP^fl/fl^-tdTomato cassette in which tdTomato is only produced upon Cre-mediated excision of the premature STOP codon^39^. The cells were cultured for two days with IL-2 and anti-CD3/CD28 beads to activate and expand the T cells and were then treated with LNP^HA^ +/-bsAb^HA-CD4^. After three days, tdTomato expression was measured in CD4+ and CD4-T cells by flow cytometry.

With SM-102-based LNP, ∼10% of CD4+ and CD4-T cells were tdTomato+, indicating no specific tropism of the particles (Fig. 3g). With SM102-based LNP^HA^ without the bsAb, there was no tdTomato+ cells, further demonstrating that the modified particle has a reduced transfection efficiency. However, when the LNP^HA^ was coupled with bsAb^HA-CD4^, CD4+ T cell transfection was made possible, with the efficiency similar to SM-102-LNP. Importantly though, there was negligible transfection of CD4-negative T cells (Fig. 3g). This further demonstrates the ability of bsAb-conjugated LNPs to mediate target cell-specific delivery of RNA.

We next evaluated *in vivo* delivery of the particles to T cells. Cre mRNA was encapsulated into SM102-based LNP and LNP^HA^, incubated with or without bsAb^HA-CD4^, and intravenous injected into Ai14 mice. After 3 days, mice were sacrificed, and splenocytes analyzed for tdTomato expression by flow cytometry. With the standard SM-102-based LNP, there was splenocyte transfection, but the majority of cells were

CD4-negative and virtually no CD4+ cells were transfected (Fig. 3h). The transfected cells were likely mononuclear phagocytic cells, such as monocytes and DCs, which are more efficiently transfected by LNPs than lymphocytes. With the standard LNP or LNP^HA^ without the bsAb, CD4+ cells were not tdTomato+, indicating they were not transfected. Instead, when mice were injected with LNP^HA^/bsAb^HA-CD4^, ∼10% of the CD4+ cells were tdTomato+ (1% of the 9% of splenocytes that were CD4+), which is a 16-fold increase over the standard particle (Fig. 3h-j). When we analyzed the CD4-negative compartment we observed reduced Cre recombination in mice treated with the LNP^HA^/bsAb^HA-CD4^ complexes suggesting a higher degree of specificity of LNP^HA^/bsAb^HA-CD4^ compared to standard LNP formulation or LNP^HA^ without bsAb (Fig. 3i, 3j).

CD5 is an endocytic receptor expressed by all T cells, which has been used as a target of antibody-targeted LNPs^18^. To see if CD5 could serve as a target for the bsAb- coupled LNP, we generated a bsAb composed of the scFV of human CD5 antibody to create bsAb^hCD5-HA^. For cell engineering, T cells are normally activated ex vivo to enhance their transduction, however this has the caveat of producing more exhausted T cells^40^. Thus, it is beneficial if LNP transfection of naïve or quiescent T cells can be enhanced. We encapsulated GFP mRNA in SM-102 LNP or LNP^HA^ with or without bsAb^hCD5-HA^, and transfected unstimulated human T cells isolated from peripheral blood. Treatment with standard LNP formulation was associated with very low levels of GFP expression, whereas with LNP^HA^/bsAb^hCD5-HA^, ∼70% of T cells became GFP+ (Fig. 3k). In addition to a higher frequency of transfected T cells, the absolute expression of GFP in the cells increased by ∼4-fold with the bsAb-coupled LNP compared to the standard LNP formulation (Fig. 3l).

These results further demonstrate that bsAb-coupled LNPs can achieve efficient cell-specific RNA delivery and reduce off-target transfection in vitro and in vivo, including enhancing transfection of unstimulated/quiescent human T cells.

### bsAb-coupled LNPs mediate comparable or enhanced transfection efficiency compared to chemically-conjugated Ab-decorated LNPs

We wanted to compare the efficiency of LNP targeted-delivery using the bsAb platform and maleimide-thiol chemistry, which is commonly used to conjugate antibodies to LNPs^14,17,22^. We encapsulate GFP mRNA in LNP^HA^ or in malemide-PEG containing LNP (LNP^MAL^). In both cases, the same lipid formulas were used and only the PEG differed. To attach an antibody to the LNP^MAL^, we first functionalized the targeting antibody (anti-mPD-L1) or control isotype-matched IgG with N-succinimidyl S-acetylthioacetate (SATA) to introduce sulfhydryl groups. The unreacted components were removed by ultra centrifugal filters, and the reactive sulfhydryl group on the antibodies were then conjugated to LNP^Mal^ through thioether conjugation chemistry by overnight incubation. For each antibody, we performed a titration of the maleimide-to-mAb ratio to identify the optimal working conditions. In parallel, we conjugated LNP^HA^ by incubating with bsAb^HA-mPDL1^.

To compare delivery efficiency, we transfected K562^mPDL1^ cells with the different formulations. Flow cytometry showed similarly high levels of targeted cell transfection by both conjugation technologies, with ∼97% of mPD-L1+ K562 cells becoming GFP+ (Fig. 4a). Notably, there was a higher degree of specificity with the bsAb/LNP^HA^, in which only ∼4% of PD-L1-negative cells were GFP+, compared with LNP^Mal-mPDL1^, which resulted in ∼37% of GFP+ PD-L1-negative cells. Similar results were obtained when we delivered Cre mRNA to K562^LoxP_GFP_mPDL1^ with LNP^HA^/bsAb^HA-PDL1^ and LNP^Mal-mPDL1^(Fig. S4a). It is unclear why the bsAb/LNP^HA^ was more specific, but this is consistent with our other data indicating that the LNP^HA^ has a reduced non-specific tropism.

**Figure 4.**
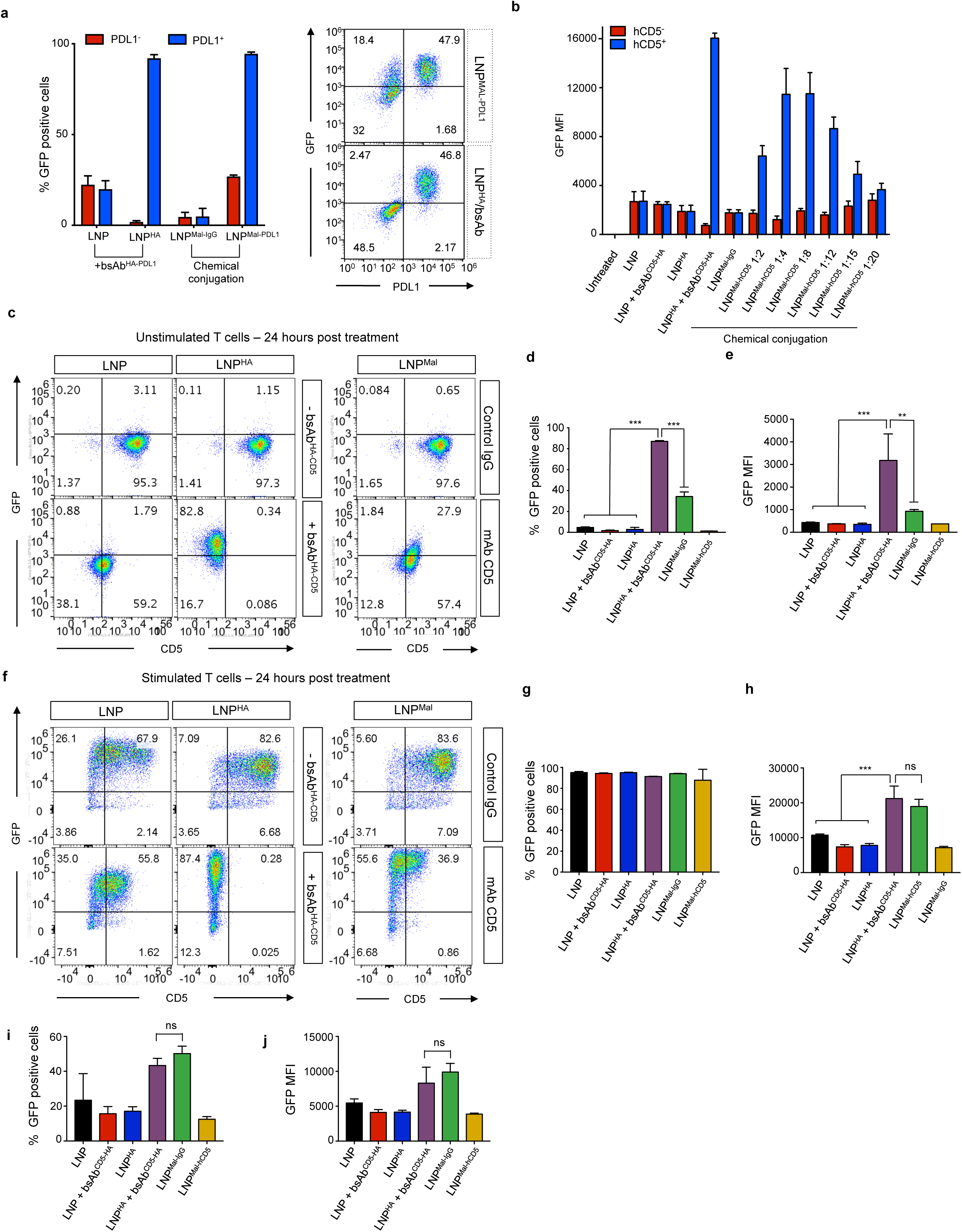
Comparison of cell transfection by LNP^HA^/bsAb and thiol-Mal-conjugated LNP. a. Flow cytometry analysis of GFP and PD-L1 expression in K562^mPDL1^ cells transfected with the indicated LNP. Graph (left) shows the mean±s.d. of GFP+ cells. Dotplots (right) are representative from the experiment. All groups used matched concentrations of LNP-RNA (10 ng). GFP expression was measured 24 hours post transfection. (n=3 per group, 3 independent experiments performed) b. Flow cytometry analysis of GFP and CD5 expression in K562^hCD5^ transfected with 10 ng GFP mRNA encapsulated in the indicated LNP. Graph shows the mean±s.d. of GFP mean florescence intensity (MFI) of GFP+ cells. (n=3 per group, 3 independent experiments performed) c. Flow cytometry analysis of GFP and CD5 expression in resting human T cells treated with 200 ng GFP RNA encapsulated in SM-102 LNP (left), SM-102 LNP^HA^ (center) with or without bsAb^CD5-HA^ or with LNP^MAL-hCD5^. Representative dotplots are shown (n=3 per group, 3 independent experiments performed). d. Graph showing the mean±s.d. of the percentage of GFP+ resting human T cells treated as in (c). One-way anova and Tukey’s multiple comparison post-test, ***p<0.0001; Data representative of 3 independent experiments with n=3 per group. e. Graph showing the mean±s.d. GFP MFI of resting human T cells treated as in (c). Statistical analyis: One-way anova and Tukey’s multiple comparison post-test, **p<0.001 ***p<0.0001 (n=3); Data representative of 3 independent experiments. f. Flow cytometry analysis of GFP and CD5 expression in activated human T cells treated with 200 ng GFP mRNA encapsulated in SM-102 LNP or SM-102 LNP^HA^ with or without bsAb^CD5-HA^, or LNP^MAL-hCD5^ or LNP^MAL-IgG^. Expression was measured 24 hours post-transfection. Data representative of 3 independent experiments (n=3 per group per experiment). g. Graph showing the mean±s.d. percentage of GFP+ activated human T cells treated as in (f). One-way anova and Tukey’s multiple comparison post-test, ***p<0.0001 (n=3); Data representative of 3 independent experiments. (n=3 per group, 3 independent experiments performed) h. Graph showing the mean±s.d. GFP MFI of activated human T cells treated as in (g). (n=3 per group, 3 independent experiments performed) i. Graph showing the mean±s.d. percent of GFP+ activated human T cells treated as in (g) at 3 days post transfection. (n=3 per group, 3 independent experiments performed) j. Graph showing the mean±s.d. GFP MFI in activated human T cells treated as in (g) at 3 days post transfection. (n=3 per group, 3 independent experiments performed)

LNP^Mal^ chemically-conjugated with anti-CD5 mAb have been used to enhance transfection of primary human T cells^18^. To see how bsAb non-chemical conjugation would compare, we encapsulated GFP mRNA in LNP^Mal^, and then conjugated with different molar ratios of maleimide:Ab. We first tested on K562 cells expressing human CD5 (hCD5) to find the optimal mal-PEG:Ab ratio. The highest transfection efficiency was achieved with LNP made with a 4:1 molar ratio of PEG^Mal^:Ab. There was reduced transfection efficiency with lower and higher PEG^Mal^:Ab ratios, indicating that insufficient and saturating doses of Ab were used in these treatment conditions respectively (Fig. 4b, Fig. S4b).

We isolated human T cells from peripheral blood, left them unstimulated, and transfected with concentration matched GFP mRNA encapsulated in the different conjugated particles (bsAb or covalent chemistry). The bsAb approached was considerably more efficient, with >80% of unstimulated T cells becoming GFP+ compared to ∼40% GFP+ T cells with the LNP^Mal-hCD5^. This was also reflected in the level of GFP in the cells, which more than doubled with the bsAb coupled particle compared to the chemically conjugated LNP (Fig. 4c-e).

This result was surprising to us as we had expected the particles to be similar in transfection efficiency. In published studies with anti-CD5 chemically-conjugated LNPs, in vitro testing was with activated T cells^41^. We thus repeated transfections using primary T cells activated with anti-CD3/CD28 magnetic beads and IL-2 for 2 days. In this setting, the same bsAb/LNP^HA^ and LNP^Mal-hCD5^ particles had similar transfection efficiencies (Fig. 4f). Though with activated T cells, even the unconjugated LNPs resulted in high transfection, albeit the anti-CD5 decorated particles (bsAb or Mal) resulted in higher GFP MFI, indicating the antibody did enhance transfection (Fig. 4g). Differences across conditions were even more evident when cells were analyzed at 4 days post transfection (Fig. 4h, 4i).

Overall, these results indicate that chemical-free conjugation of LNPs using a bsAb is as efficient as the covalently conjugated particles, and that the bsAb approach can even enhance LNP transfection of unstimulated T cells.

### A tethering bsAb enables non-chemical conjugation of off-the-shelf antibodies with LNPs

LNP^HA^/bsAb complexes enable targeted LNP cell transfection without chemical conjugation of the LNP but require the production of a new bsAb for each molecular target. To minimize the need for bsAb development, we hypothesized that we could modify the non-LNP binding portion of the bsAb to recognize the constant region of an antibody, and use this bsAb to tether the LNP to any antibody with the corresponding constant region (Fig. 5a). To this end, we generated a bsAb with the scFV for HA and a second scFv that binds the constant region of rat IgG2b to create bsAb^HA-IgG2b^.

**Figure 5.**
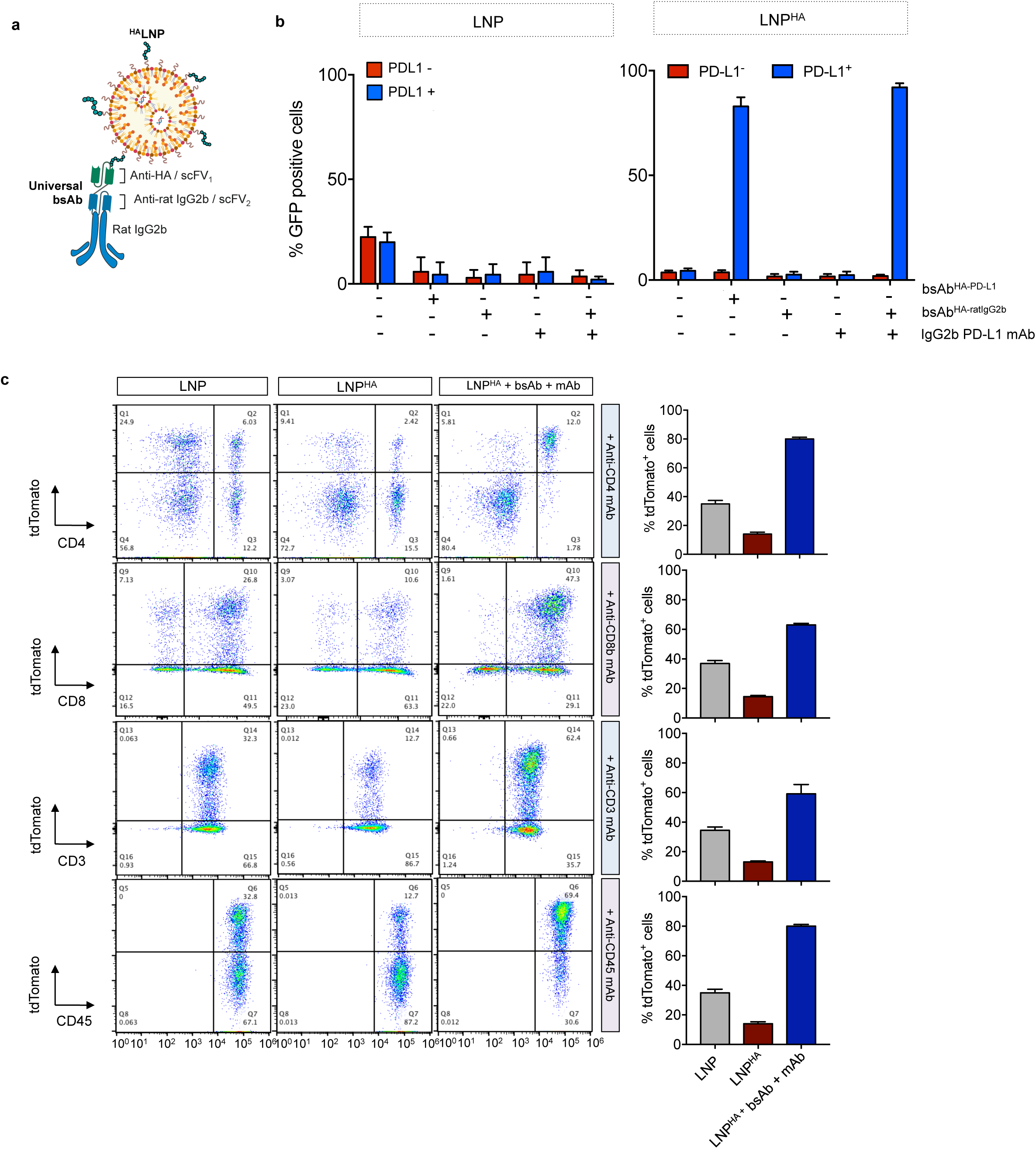
Tether bsAb platform redirect LNP^HA^ towards target cell type in presence of monoclonal antibody. a. Schematic depicting the bsAb^HA-ratIgG2b^ tethering antibody binding to LNP^HA^ particles and a monoclonal antibody with the recognized constant region (Rat IgG2b). b. Flow cytometry analysis of GFP expression in K562^mPDL1^ cells transfected with 10 ng of GFP mRNA encapsulated in SM-102 LNP or SM-102 LNP^HA^ with or without bsAb^HA/PD-L1^ or with bsAb^HA/ratIgG2b^ with or without PD-L1 monoclonal antibody. GFP expression was measured 24 hours post transfection. Graphs show the mean±s.d. of the percentage of GFP+ cells. c. Representative dotplots and graphs of total splenocytes treated with Cre mRNA encapsulated in LNP or LNP^HA^ with or without bsAb^HA-IgG2b^ and the indicated monoclonal antibody. Analysis was performed 48 hours after LNP treatment. Data representative of 3 independent experiments.

To test the bsAb^HA-IgG2b^ tethering system, we selected a rat anti-murine PD-L1 antibody with an IgG2b constant region. We encapsulated GFP mRNA in LNP^HA^ and incubated with either bsAb^HA-PDL1^ or with bsAb^HA-IgG2b^ plus anti-PD-L1 (tethered LNP). K562^mPDL1^ were treated with equal concentrations of the particles, as well as control particles that did not include one or another antibody, or the parent particle formulation that did not incorporate PEG^HA^. Transfection with the tethered LNP resulted in ∼90% of the mPD-L1+ K562 cells being GFP+ (Fig. 5b). This was similar in efficiency to LNP^HA^/bsAb^HA-PDL1^. Importantly, this was specific, as LNP^HA^/bsAb^HA-IgG2b^ alone (i.e. without anti-PD-L1 tethered) resulted in <5% of GFP+ PD-L1+ cells. The tethered LNP was also far more efficient than the standard LNP at transfecting the K562 cells (Fig. 5b). Similar results were obtained when we delivered Cre mRNA to K562^LoxP_GFP_mPDL1^ cells using the tethered LNP (Fig. S5a). These data indicates that the dual component bsAb tethering system is as efficient as the single bsAb in targeting LNP-RNA to PD- L1+ cells.

To further assess the approach, we took LNP^HA^/bsAb^HA-IgG2b^ particles (GFP mRNA encapsulated) and incubated with or without a rat anti-CD20 antibody that has an IgG2b constant region, in order to tether the anti-CD20 to the LNP. We then transfected BCL leukemic B cells, which express CD20. At the tested dose, we observed very low level of transfection with standard SM-102 LNP as well as with SM-102 LNP^HA^ in absence of the bsAb (Fig. S5b). However, with LNPs in which anti-CD20 was tethered via bsAb^HA-IgG2b^, ∼60% of BCL1 cells became GFP+ (Fig. S5b). This demonstrated that the bsAb^HA-IgG2b^ was able to tether the anti-CD20 antibody to the LNP and enhance transfection.

We reasoned that the bsAb^HA-IgG2b^ would enable us to easily test many different antibodies for targeted LNP delivery of RNA. We encapsulated Cre mRNA in LNP or LNP^HA^. An aliquot of LNP^HA^ was incubated with bsAb^HA-IgG2b^. All three formulations (LNP, LNP^HA^, LNP^HA^/bsAb^HA-IgG2b^) were then incubated with anti-CD4, anti-CD8, anti-CD3, and anti-CD45 monoclonal antibodies with rat IgG2b constant regions. As all four antibodies target molecules expressed by T cells, we isolated primary murine T cells from the spleens of Ai14 mice and transfected them with each formulation. In all cases, the bsAb^HA-IgG2b^ was able to tether the monoclonal antibody to the LNP and mediate enhanced delivery to the T cells expressing the targeted molecule, as indicated by GFP expression (Fig. 5c).

These data demonstrate that bsAb^HA-IgG2b^ can be used with different rat IgG2b monoclonal antibodies to direct RNA-LNP to cells of interest and highlight the flexibility of our conjugation approach in generating antibody-directed LNPs.

## DISCUSSION

Here we employed LNP surface engineering and the use of a bsAb to establish a simple and efficient technology to enhance LNP-mediated RNA delivery to desired cell populations. Assembly of the bsAb/LNP complexes occurs in physiological conditions without the need for chemical coupling reagents or additional purification steps and with unidirectional orientation of the bsAb on the LNP surface. This can be performed by any lab, and, using the tethering bsAb, with off-the-shelf antibodies, providing a broadly applicable platform for targeted LNP delivery.

Direct antibody conjugation of the LNP allows efficient cell-specific targeting and is a technology that can greatly expand the applications of LNP technology by enabling more efficient delivery to selected cells in vitro or in vivo^24^. However, the nature of the chemical conjugation method makes the technology difficult to standardize and manufacture at higher scale. There can be batch-to-batch variations^23^, which may come from the random orientation that the Ab conjugates to the particle’s surface, owing to multiple thiol groups on the antibody. There is also the need for additional purification steps to remove cross-linking and catalytic reagents.

The bsAb approach to LNP conjugation avoids the need for chemical conjugation of the antibody to the LNP, but still provides a means to decorate the particle with a targeting antibody. Importantly, the approach was effective across all FDA-approved LNP formulations, on primary human cells of clinical relevance, and for targeted in vivo delivery. LNP^HA^ can be produced at scale and retain the same biophysical characteristics of standard LNP formulations, including PDI, charge and mRNA encapsulation efficiency. Interestingly, addition of PEG^HA^ to the LNP resulted in a general decrease in cell transfection efficiency in absence of bsAb. This had the important benefit of increasing specificity of LNP targeting by reducing transfection of non-target-expressing cells, while target-expressing cell transfection was enhanced. In contrast, we found that antibody-conjugated LNPs generated by chemical conjugation still resulted in significant non-target cell transfection. It is not exactly clear why the LNP^HA^ particle had this property of low off-target transfection, especially given that just 15% of the LNP’s PEG was PEG^HA^. It could be that the HA peptide affects non-receptor-mediated endosomal escape or particle binding to serum proteins.

We generated and validated multiple bsAb and demonstrated that this technology can be used across different cellular targets. This included clinically relevant targets such as PD-L1 and CD5, but also the SunTag^31^, which can be fused to different cell surface proteins and used as a ‘tool’ target, potentially even for screening the targetability of different receptors for enhancing LNP uptake. As an even more flexible approach, we established a bsAb that can bind off-the-shelf monoclonal antibodies. This obviates the need to generate a new bsAb for each target, and greatly broadens the potential utility of the approach, especially for labs that do not have expertise in chemically conjugating LNPs or generating bsAb. The ‘bridging’ bsAb we generated recognizes the constant region of rat IgG2b and can thus tether an LNP to any antibody with this domain. A similar bsAb could be made to recognize other antibody constant regions, which would make even more antibodies available for LNP targeting. We envision this being particularly useful to screen multiple antibodies and/or surface receptors for their capability to mediate cell-specific LNP transfection.

We used the HA peptide as the docking target of the bsAb because HA is a short linear epitope and the antibody is known to bind with high affinity. However, we expect that other short linear epitopes could be used in place of HA. There have also been reports that PEG itself could be targeted by a bsAb^42^ ^43^. This strategy eliminates the addition of the HA. However, a shortcoming of this approach is that it is harder to titrate the amount of antibody decoration, since all the PEG can be bound by the antibody. In addition, it was reported that binding of the PEG bsAb on the LNP surface induced LNP destabilization and particle aggregation^44^. We did not observe any major difference in transfection efficiency when we either pre-coupled the LNP^HA^ and bsAb or when we provided the two components simultaneously, indicating the bsAb binding to HA does not destabilize the LNP. As noted above, the LNP^HA^ also enhanced specificity by reducing transfection of non-targeted cells. Thus, there is utility for the inclusion of a separate dock that is distinct from the PEG. It is worth noting that the HA epitope lies within a non-immunogenic region of the native HA protein^45^. Even if antibodies do develop against the HA, as they can with PEG^46^, we expect the epitope to be occluded by the bsAb, but alternative epitopes could also be utilized.

Antibody-conjugated LNPs have the capacity to enable cell-targeted delivery, which can be critical when the RNA cargo encodes a protein with toxic potential, such as a CAR^18^ or immunostimulatory cytokine like IL-12^47^. Additional to specificity, Ab-conjugated LNPs have utility for enhancing transfection of target expressing cells, even in vitro. For example, anti-CD117 conjugated LNPs enhanced ex vivo HSC transfection by ∼3-fold, compared to parent particles^21^. We found this as well with our CD3, CD4, and CD5 bsAb targeted LNPs. Of note, while transfection of activated T cells was obtained with both bsAb and Ab-conjugated particles, transfection of unstimulated human T cells was achieved exclusively with LNP^HA^/bsAb^HA-CD5^. To our knowledge, this is one of the most efficient demonstrations of LNP transfection of unstimulated human T cells. This is impressive, as normally T cells are activated for transfection or even transduction with lentiviral and retroviral vectors, and this can have biological consequences, such as accelerating exhaustion^48^. Use of the LNP^HA^/bsAb^HA-CD5^ may enable genetic manipulation of unstimulated T cells for more potent control of polarization.

Antibody-mediated delivery of LNPs has been established as an effective means to enhance RNA delivery to specific cell types^49^. Through the use of bsAb and inclusion of a docking peptide, functionalizing LNPs with an antibody targeting moiety is made simple and efficient, while maintaining particle integrity. This should expand the useability and applicability of the technology, while enabling enhanced LNP-RNA delivery to cells of interest.

## ACKNOWLEDGMENTS

The authors thank members of the Brown and Dong labs for their helpful comments. AM was supported by NIH T33AI078892. CF was supported by Cancer Research Institute Irvington Fellowship (CRI4641), BDB was supported by NIH R01DK138025 and R01AI104848 and the Applebaum Foundation.

## AUTHOR CONTRIBUTIONS

Conceptualization, A.A., B.D.B. A.J.P.T.; methodology, A.A.,; investigation, M.P., Z.Y., P.S., A.M., J.M.F., M.S., G.M., C.F.,; writing – original draft, A.A., B.D.B.; writing –review and editing, A.A., J.B., Y.D.,B.D.B; supervision, B.D.B.; funding: B.D.B.

## MATERIALS & METHODS

### Lipids

SM-102 (CAS# 2089251-47-6), ALC-0315 (CAS#: 2036272-55-4) and ALC-0159 (CAS#: 1849616-42-7) were purchased from MedKoo Biosciences. (6Z,9Z,28Z,31Z)- Heptatriaconta-6,9,28,31-tetraen-19-yl 4-(dimethylamino)butanoate (D-Lin-MC3-DMA) (Cat#1224606-06-7) was purchased from Broad Pharm. Cholesterol (CAS# 57-88-5) was purchased from Millipore Sigma. 1,2-dimyristoyl-rac-glycero-3- methypolyoxyethylene (DMG-PEG2000) was purchased from NOF America Corporation. 1,2-disteraroyl-sn-glycero-3-phosphocholine (DSPC) (SKU# 850365P- 25mg), 1,2-dioleoyl-sn-glycero-3-phosphoethanolamine 18:1 (Δ9-Cis) PE (DOPE) and 1,2-distearoyl-sn-glycero-3-phosphoethanolamine-N-[maleimide(polyethylene glycol)- 2000] (DSPE-PEG2000 Maleimide, PEG^Mal^) (SKU# 880126P-10mg), were purchased from Avanti Polar Lipids. C-GGGYPDDVPDYA DSPE-PEG2000-Mal conjugation on Cys (PEG^HA^) (Cat# 37182) was purchased from LifeTein, LLC.

### Chemicals

Pierce® SATA (N-succinimidyl S-acetylthioacetate) (Cat#26102) and Pierce® Hydroxylamine-HCl (Cat#26103) were purchased from ThermoFisher Scientific. Buffered formaldehyde (4%, J60401-AK) was purchased from Thermo Fischer Scientific. 0.1x TE was obtained from Maxi Qiagen kit #12362 purchased from Qiagen. 0.25M CaCl2 (Cat#C7902–1KG) was purchased from Sigma Aldrich. 2x HBS was prepared in house. For 500mL: 1M HEPES (50ml Corning #25–060-Cl), 2M NaCl (70.25ml Fisher Bioreagents #BP358–1); 0.5M Na2HPO4 (1.5ml # BP332–500), 378.25ml Tissue Culture Tested Water (Corning #46–000-CV), 5M NaOH to pH. Quant-it™ RiboGreen RNA Assay Kit and RiboGreen RNA Reagent was purchased from ThermoFisher Scientific (Cat# R11491).

### Cells and cell culture conditions

293T cells (embryonic kidney; human) and ID8 Brca1-deficient cells (ovarian adenocarcinoma; C57BL/6 mice origin) were grown in IMDM with 10% heat-inactivated fetal bovine serum (FBS), 100 U/ml penicillin/streptomycin and 2 mM L-Glutamin. 293T cells were purchased from ATCC (#CRL-3216). ID8 Brca1-deficient cell line was kindly provided by Jean J. Zhao (Harvard). B16-F10 cells (C57BL/6 mouse; ATCC #CRL-6475) were cultured in Dulbecco’s modified Eagle medium (DMEM) containing heat-inactivated 10% FBS, 100 U/ml penicillin/streptomycin and 2 mM L-Glutamin. K562 (human; ATCC #CCL-243^TM^) and BCL-1 (BALB/c mice; ATCC #TIB-197 ™) were cultured in Roswell Park Memorial Institute (RPMI) medium containing 10% heat-inactivated FBS, 100 U/ml penicillin/streptomycin and 2 mM L-Glutamin. The KC pancreatic cancer cell line was kindly provided by Dr. Dieter Saur (Technical University of Munich) and cultured in DMEM containing heat-inactivated 10% FBS, 100 U/ml penicillin/streptomycin and 2 mM L-Glutamin. MEL cell line was cultured in RPMI containing heat-inactivated 10% FBS, 100 U/ml penicillin/streptomycin and 2 mM L-Glutamin.

Ai14 splenocytes were cultured in RPMI 10% FBS with 500 U ml−1 IL-2, 2-Mercaptoethanol (Thermo Fisher Scientific, Cat # 21985023), and activated with Gibco™ Dynabeads™ Mouse T-Activator CD3/CD28 (Fisher Scientific, Cat # 11452-D) for 2 days prior LNP treatment. Human primary T lymphocytes were isolated from peripheral blood mononuclear cells of healthy donors by leukapheresis and Ficoll-Hypaque gradient separation. Cells were enriched to purity using EasySep™ Human T Cell Isolation Kit (Cat# 17951) according to manufacturer instructions. Unstimulated T cells were cultured at a concentration of 1 × 10^6^ cells/ml in RPMI supplemented with penicillin, streptomycin, 10% FBS and and 2-Mercaptoethanol. Stimulated T cells were cultured in presence of recombinant human IL-2 500 U ml−1 (Peprotech) and Gibco™ Dynabeads™ Human T-Activator CD3/CD28 (Fisher Scientific, Cat # 11161-D) for 2 days before LNP treatment.

### Mouse Strains

In vivo studies were performed using C57BL/6 (JAX# 000664) and Ai14 (JAX# 007914) mice obtained from Jackson laboratories and housed in the Mount Sinai vivarium during use. All mouse experiments were carried out under institutional IACUC approval. All mice were randomized before experimentation.

### Lentiviral vector production

Lentiviral vector production was performed as previously described in detail^36^. Briefly, HEK293T cells were seeded in 15 cm tissue culture plates (Thermo Scientific Nunclon #168381) 24 hours prior to achieve an approximate cell density of 70% at the time of transfection. Transfection was carried out using the calcium-phosphate method. Reporter plasmid constructs were mixed with packaging plasmids (Gag, Rev, Pol) and suspended in 0.1x TE and 0.25M CaCl2; one volume of 2x HBS was added in a dropwise fashion while continually vortexing, and the resulting solution was immediately added onto HEK293T cells and allowed to sit overnight. IMDM medium was replaced the next morning and supernatants collected and 0.2 µm-filtered 24–30h after that. Lentivirus supernatant aliquots were stored at −80°C until use.

### Bispecific antibody (bsAb) production and purification

All bsAbs were produced using the Expi293T^TM^ expression Kit (Gibco, Cat #A14635) according to the manufacturer instructions. For all transfections a ratio of 1.0 ug plasmid DNA/mL of transfection culture was used. Transfections were performed on Expi293 cells in a final volume of 30 mL in 125 mL non-baffled flasks (Greiner Bio-One, Cat. 679501). Cultures were maintained on a CO_2_ resistant shaker with an orbit of 19 mm (Thermo Fisher, Cat. 88881101). Expi293F cells were cultured in Expi293 Expression Medium (Gibco, Cat. A1435101) supplemented with 1% penicillin-streptomycin (Gibco, 15140-122) in a humidified 8% CO2 incubator at 37°C and 125 RPM. Supernatant containing bsAb was harvested 4 days post-transfection, centrifuged at 3000 x g for 25 minutes at 4°C and filtered through a 0.2 µM PES filter (Thermo Scientific, Cat. 564-0020).

bsAb possessing a His tag were purified using Ni-NTA agarose (QIAGEN, Cat. 30210) with gravity flow columns (Marvelgent Biosciences Inc., Cat 12-0278-050), as described^50^. 2.5 mL of Ni-NTA agarose were used for every 30 ml supernatant. Briefly, Ni-NTA agarose was washed 3 times with molecular biology grade water (Corning, Cat. 46-000-CV). The supernatant from the transfection was incubated with the Ni-NTA agarose in slow agitation for at least 60 minutes at 4°C.1 L of binding buffer was prepared by mixing 25 mL of 1 M Tris-HCL with a pH of 7.5 (Fisher Bioreagents, Cat. BP1757-500), 200 mL of 2.5 M NaCl (Fisher Bioreagents, Cat. BP358-1), 50 mL of glycerol (Research Products International, Cat. G22025-0.5), 70 µL of 2-mercaptoethanol (Gibco, Cat. 21985023), 1 mL of 5 M imidazole (Fisher Bioreagents, O3196-500), and 723.93 mL of molecular biology grade water. 1 L of 1 M imidazole elution buffer was prepared by mixing 25 mL of 1 M Tris-HCL with a pH of 7.5, 200 mL of 2.5 M NaCl, 50 mL of glycerol, 70 µL of 2-mercaptoethanol, 200 mL of 5 M imidazole, and 524.93 mL of molecular biology grade water. All solid reagents had been resuspended in molecular biology grade water to reach the indicated concentration. The binding and elution buffers were mixed to create elution buffers with varying imidazole concentrations as follows: 10 mM, 15 mM, 50 mM, 250 mM, and 500 mM. A gravity flow column was equilibrated with 20 mL of binding buffer. The supernatant/agarose mixture was passed through the column and the flow through was collected. The column was then washed twice with 10 mL of binding buffer; 10 mL each of 10 mM and 15 mM imidazole elution buffers; and 6 mL each of 250 mM, 500 mM, and 1 M imidazole elution buffers. Each wash fraction was collected in separate conical tubes and was analyzed by loading a sample onto an SDS-PAGE gel. Fractions that contained the bsAb were combined and concentrated to a final volume of 1 mL by centrifuging at 6,000 x g at 4°C in a Vivaspin 20 Centrifugal Concentrator (Sartorius, Cat. VS2002). The concentrated bsAb was placed in an equilibrated dialysis cassette (Thermo Scientific, Cat. A52966) and dialyzed in 500 mL of PBS (Corning, Cat. 21-040-CV) overnight at 4°C. Dialyzed bsAb were then quantified using a NanoDrop 2000 Spectrophotometer (Thermo Scientific) and stored in 10% glycerol at-80°C.

### RNA template cloning

Templates for in vitro transcription (IVT) were generated by restriction cloning using a T7 promoter sequence containing 5’UTR, an open reading frame, and 3’UTR. Our construct employed NASAR UTRs for enhanced gene expression^51^. The addition of a 120-nulcleotide poly-A tail to the mRNAs was encoded on the PCR template with a reverse primer containing a 120-nucleotide poly-T. To generate IVT template from plasmid, we amplified DNA with Q5® High-Fidelity Master Mix (NEB, M0492). Following PCR, RNA was incubated at 37°C for 30 minutes with DpnI (NEB, R0176) to remove plasmid contaminants and PCR product was cleaned up with QIAquick® PCR Purification Kit (Qiagen). Concentration was measured using the NanoDrop™ 2000 (Thermo Scientific) for application in IVT.

### Synthesis and purification of IVT generated modified mRNA

All mRNAs were synthesized using HiScribe® T7 High Yield RNA Synthesis Kit (NEB). UTP was fully substituted with N1m (N1-methylpseudouridine-5′-triphosphate) (Crystal Chem Cat# 13946). mRNA was capped by inclusion of 1:4 premix of CleanCap® reagent AG (TriLink) and capping occurred co-transcriptionally. The reaction mixtures were incubated at 37°C for 6 hours and further incubated at 37°C for 30 min in the presence of TURBO DNaseI (Thermo Scientific). RNA products were purified with Monarch® RNA Cleanup Kit (NEB, T2050). Concentration of RNA was measured using the NanoDrop™ 2000 (Thermo Scientific) and species purity and size was assessed on a denaturing RNA gel.

### Lipid nanoparticle formulation and modified mRNA encapsulation

mRNA encapsulation in LNP was performed using the Ignite® microfluidic mixing device (Precisions Nanosystems). SM-102-baed particles^34^ were formulated with the helper lipid DSPC, cholesterol, DMG-PEG2000 (molar ratio 50:10:38.5:1.5), and mRNA dissolved in a citrate buffer. In SM-102 LNP^HA^, 15% of total PEG was substituted with PEG^HA^. ALC-0315-based LNPs were formulated with ALC-0315, DOPE, cholesterol and ALC-0159 PEG at molar ratio 46.3:9.4:42.7:1.6. D-Lin-MC3-DMA-based LNP^52^ were formulated with D-Lin-MC3-DMA, DSPC, cholesterol, DMG-PEG2000 at molar ratio 50:10:38.5:1.5. LNP^HA^ variant of these particles were also modified to include 15% PEG^HA^. After formulation, the freshly formed RNA-LNPs were dialyzed overnight against PBS buffer using Slide-A-Lyzer dialysis cassettes (3.5K MWCO, Life Technologies) and subsequently concentrated in Amicon® Ultra Centrifugal Filters (10 kDa MWCO, Sigma) to desired concentration. Particle size and zeta potential of LNPs were measured using a Zetasizer Advance (Malvern Panalytical) at a scattering angle of 173° and a temperature of 25°C. All formulations were within the following parameters: 60-100 nm average size, >90% encapsulation, polydispersity <0.2.

Encapsulation efficiency of LNPs was determined using Quant-it™ RiboGreen RNA Assay Kit according to manufacturer protocols. The concentration of LNP-mRNA was defined by the amount of mRNA encapsulated in the LNP. In general, we prepared LNP-mRNA to have a final concentration of mRNA of 0.1 mg/ml.

### Generation of LNP^HA^/bsAb complex

mRNA-loaded LNP^HA^ and the corresponding bsAb were either pre-incubated together for 15 minutes or simultaneously added to the cell culture at a 1:4 weight ratio [e.g. for 1 µg LNP-mRNA (i.e. 1 µg RNA) we incubated with 4 µg of bsAb]. For in vivo studies, LNP^HA^ and bsAb were pre-incubated for 15 minutes before injection at a 1:1 weight ratio (e.g. for 1 µg of LNP-mRNA we incubated with 1 µg of bsAb).

### Generation of LNP^Mal-Ab^

LNP^Mal^ was generated as previously described^22^. Briefly, 10% of total PEG in SM-102 LNP was substituted with PEG^Mal^. Targeting antibodies or control isotype-matched IgG was functionalized with SATA to introduce sulfhydryl groups allowing conjugation to maleimide^14^. SATA was deprotected using 0.5 M hydroxylamine followed by purification by removing the unreacted components by Amicon® Ultra Centrifugal Filters (10 kDa MWCO, Sigma). The reactive sulfhydryl group on the antibody was then conjugated to LNP^Mal^ through thioether conjugation chemistry by overnight incubation at 4°C.

### In vitro cell transfection

K562 and BCL-1 cells were plated at 10^5^ cells/well in a clear, flat-bottom 96-well plate. Cells were treated with 10ng/well LNP-mRNA and 40 ng/well bsAb, if not otherwise specified. 293T cells were plated at 10^5^ cells/well in 24 well plate. Cells were treated with 1ng/well LNP-mRNA and 4 ng/well bsAb. Isolated murine T cells were plated at 2×10^5^ cells/well in a clear, flat-bottom 96-well plate and treated with 50 ng/well LNP-mRNA and 200 ng/well bsAb if not otherwise specified. Isolated human T cells were plated at 2×10^5^ cells/well in a clear, flat-bottom 96-well plate. For both unstimulated and stimulated T cells, cells were treated with 200ng/well LNP-mRNA and 800ng/well bsAb.

### Animal experimental

For in vivo experiments, Ai14 mice were i.v injected with 1 mg/kg Cre mRNA encapsulated in LNP, which was pre-incubated with bsAb^HA-CD4^ at ratio 1:1 (weight/weight). At experimental endpoints, mice were sacrificed, and spleens were collected for flow cytometry analysis.

To generate disseminated ovarian tumors, 5×10^6^ ID8 cells were injected intraperitoneally (i.p.) into 8-10 weeks old female C57BL/6J mice. At 14 days post-tumor injection, mice were i.p. injected with 1mg/kg Cre mRNA encapsulated in LNP^HA^ plus or minus bsAb. Mice were euthanized upon ascites development and tumor tissue from the omentum was collected and processed for flow cytometry.

For the melanoma model, 10^5^ B16F10 melanoma cellswere subcutaneously (s.c.) injected into-10 weeks old female C57BL/6 mice. Mice were randomly grouped and when the tumor reached ∼50 mm^3,^ (∼1 week), 10 ug Cre mRNA encapsulated in LNP^HA^ were injected intratumorally plus or minus bsAb. At 2 days after LNP-mRNA injection, tumor tissue was collected and processed for flow cytometry.

For the orthotopic pancreatic cancer model, tumor cells derived from Kras^LSL-G12D/+;^Ptf1a^Cre/+^ (KC) mice^53^ were used. In brief, 5×10^4^ were orthotopically grafted into the pancreas of wildtype C57BL/6J mice, as previously^35^. One week after tumor injection mice were i.p. injected with 1 mg/kg Cre mRNA encapsulated in LNP or LNP^HA^ plus/minus bsAb. At 2 days after LNP injection, mice were euthanized and tumor tissue collected and processed for flow cytometry as described in the “Flow Cytometry” section.

### Flow cytometry

For flow cytometry analysis of ID8 tumors in the omentum and B16F10 melanomas, the tumor-bearing tissue was collected from mice and chopped into small pieces with scissors in digest media containing collagenase and DNAase (Sigma Aldrich) and then incubated in the same media with 500 RPM lateral shaking at 37°C for 45 min. The single cell suspension was obtained by passaging the digested tumors through a 70 µm filter and washing with PBS. Cells were centrifuged at 350g for 5 minutes at 4°C and cell pellets were resuspended with ACK lysis buffer (Life Technologies) to lyse red blood cells at room temperature for 10 minure and washed with cold PBS. Flow cytometry samples were acquired with BD LSRFortessa II (BD Biosciences) and cell sorting was performed with BD FACSAria III Sorter (BD Biosciences). Pancreatic tumor were chopped into small pieces and processed, as previously described^35^.

For analysis of tdTomato expression in T cells from Ai14 mice injected with bsAb^HA-CD4^/LNP, spleens were collected and minced using a sterile blade, homogenized by smashing, and filtered through a 70-µm cell strainer. The tissue solution was washed once with PBS and pelleted by centrifuging for 5 minutes at 350g. The supernatant was removed, and the cell pellet was resuspended in 1.5mL of 1x RBC lysis buffer. After incubation, 15 mL of PBS was added to stop red blood cell lysis. The solution was centrifuged again at 350g for 5 minute to obtain a cell pellet. The single cells were suspended in a cell staining buffer and added to flow tubes that contained antibodies (total volume 100 µL). The spleen cells were analyzed using a LSRFortessa X-20 (BD Biosciences) for tdTomato.

For analysis or sorting of cells in vitro, adherent cells were detached with 0.05% trypsin-EDTA and resuspended in the flow buffer (PBS with 2% BSA).

### Antibodies

Antibodies were purchased from Biolegend or Invitrogen as specified in the chart below.

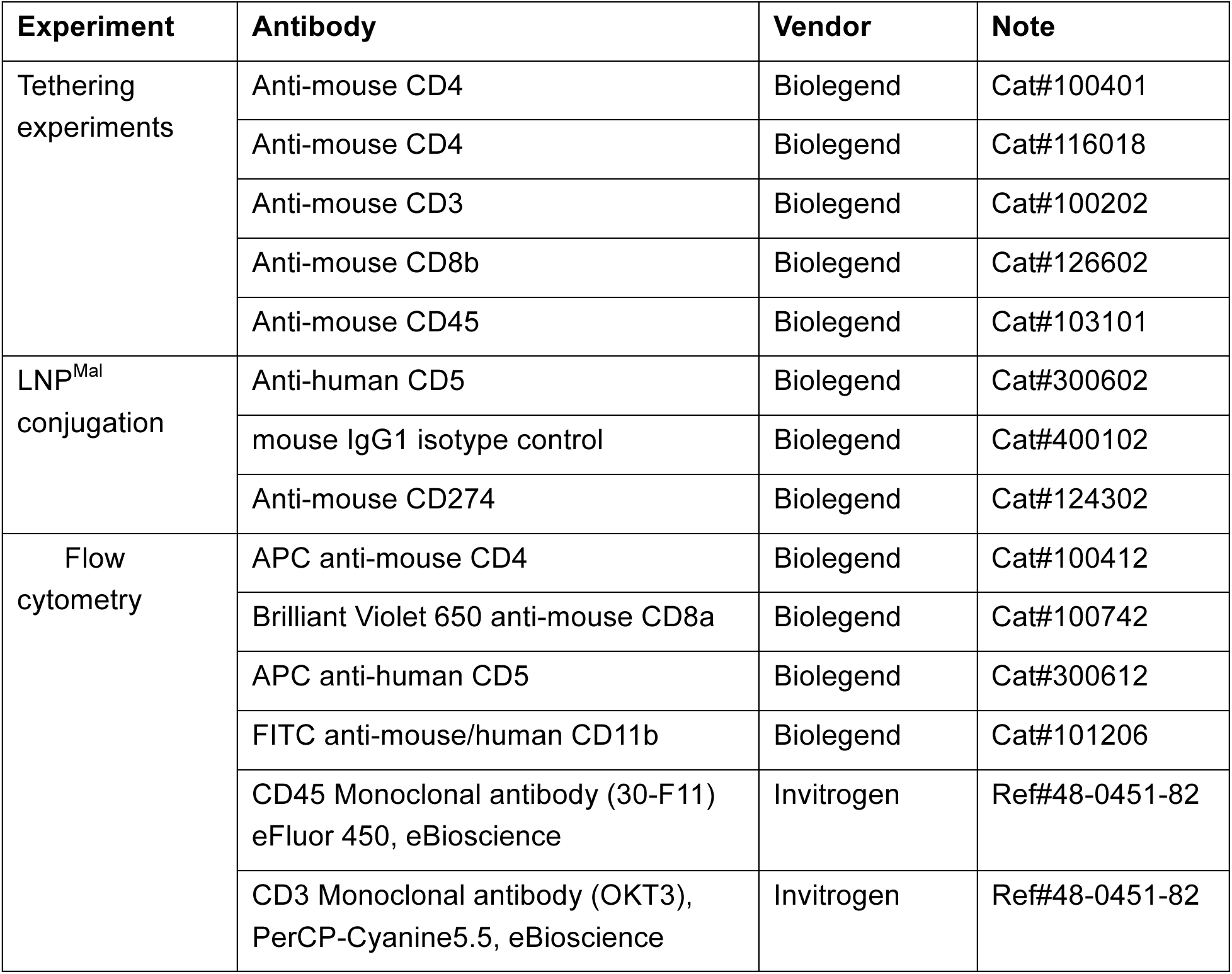
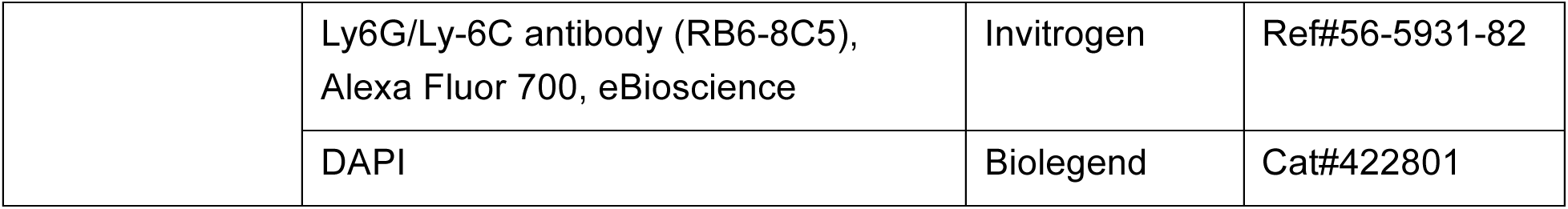

### Quantification and statistical analysis

Statistical values including the number of replicates and statistical significance are reported in the figure or figure legends when appropriate. For the majority of in vivo experiments, the experiments were repeated at least two separate times with different cohorts of mice, synthesized RNA, and encapsulated and quantified RNA-LNP. Statistical analysis was performed using Microsoft Excel or GraphPad Prism 8 (GraphPad Software Inc). Flow cytometry analysis was performed using FlowJo software. The levels of significant (unpaired two-tailed student’s t-test, one-way and two-way anova) are denoted as *p<0.05, **p<0.01, ***p<0.001 and ****p<0.001.

