## Supplementary Figures for "Bispecific antibody targeting of lipid nanoparticles"

### Supplementary Figure 1.

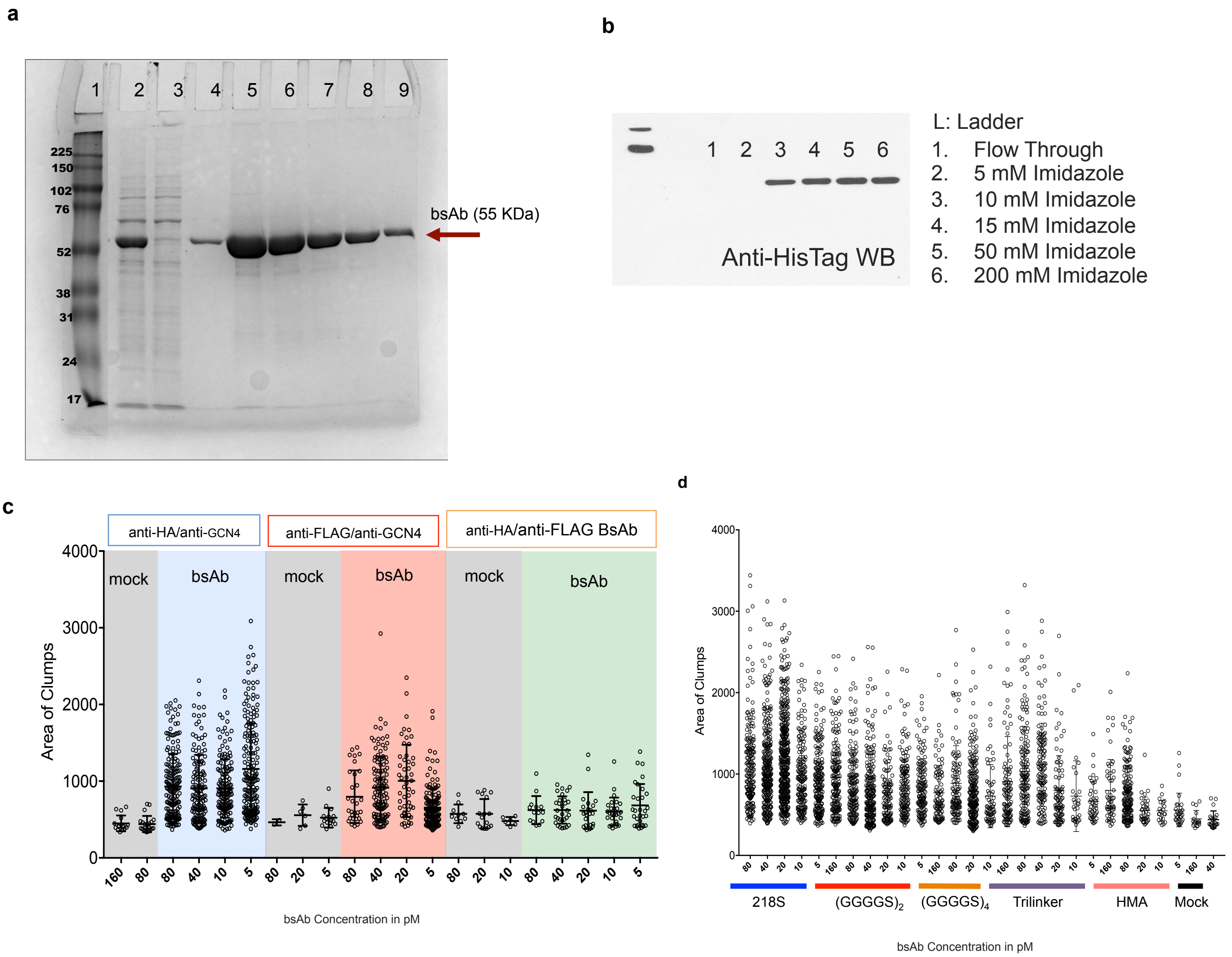

**Supplementary Figure 1. Production and purification of bsAb$^{HA\text{-}Suntag}$.**

a. SDS-PAGE analysis of the purified His6-bsAb$^{HA\text{-}Suntag}$ 1: ladder; 2: Input 3: Flow through; 4: 10mM Imidazole wash: 5-7: 50mM Imidazole elution fractions; 8-9: 250 mM Imidazole elution fractions.

b. Western blot analysis of purified bsAb$^{HA\text{-}Suntag}$

c. Histogram showing  number and area of clumps cross a dose curve induced by bsAb$^{HA\text{-}Suntag}$, bsAb$^{FLAG\text{-}Suntag}$ and bsAb$^{HA\text{-}FLAG}$ measured through by Amnis® Image-Stream analysis.

d. Histogram showing number and area of clumps cross a dose curve induced by bsAb$^{HA\text{-}Suntag}$ with different protein linker measured through by Amnis® Image-Stream analysis. Trilinker: 218s-(GGGGS)x2-HMA HMA:Human Aldolase protein linker

### Supplementary Figure 2.

a

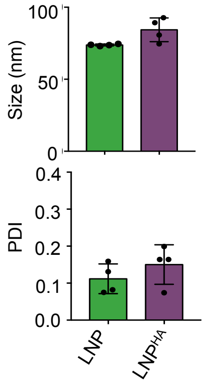


b

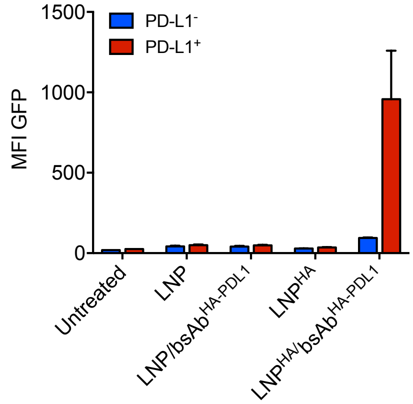


c

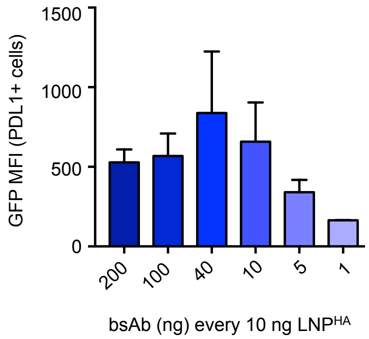


d

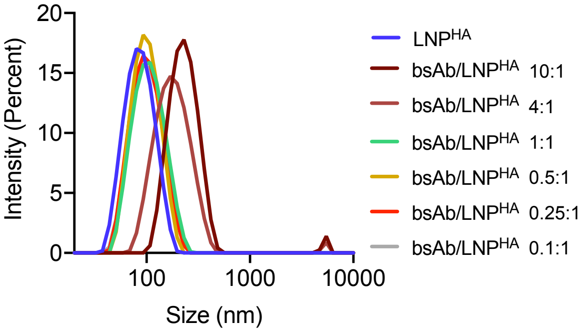


| LNP | SIZE | PDI |
|---|---|---|
| LNP$^{HA}$ | 82.93 ± 2.41 | 0.15 ± 0.04 |
| bsAb/LNP$^{HA}$ 10:1 | 245.3 ± 14.77 | 0.28 ± 0.06 |
| bsAb/LNP$^{HA}$ 4:1 | 179.4 ± 4.54 | 0.23 ± 0.04 |
| bsAb/LNP$^{HA}$ 1:1 | 97.23 ± 0.84 | 0.13 ± 0.02 |
| bsAb/LNP$^{HA}$ 0.25:1 | 93.93 ± 1.05 | 0.14 ± 0.06 |
| bsAb/LNP$^{HA}$ 0.1:1 | 93.01 ± 1.86 | 0.23 ± 0.02 |

e

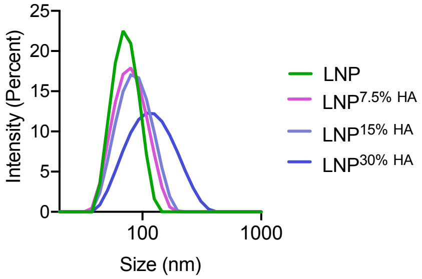


| LNP | SIZE | PDI |
|---|---|---|
| LNP | 69.66 ± 0.4 | 0.04 ± 0.01 |
| LNP$^{HA}$ 7.5% HA | 74.91 ± 0.37 | 0.08 ± 0.029 |
| LNP$^{HA}$ 15% HA | 81.32 ± 0.59 | 0.102 ± 0.04 |
| LNP$^{HA}$ 30% HA | 108.7 ± 1.47 | 0.17 ± .0.02 |

f

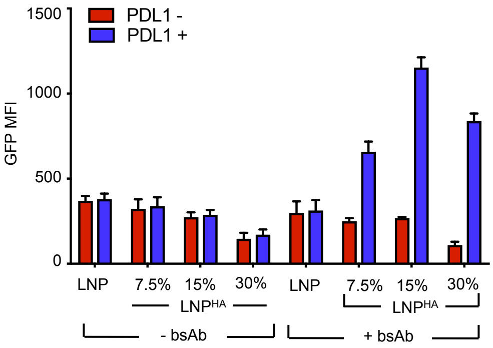

**Supplementary Figure 2. LNP$^{HA}$/bsAb complex characterization**

a. Histograms showing PDI (left) and size (right) of of LNP (green) and LNP$^{HA}$ (purple). (n=4 per group, 10 independent experiments performed)

b. Histograms showing the MFI of the GFP positive K562$^{PD-L1/Suntag}$ cells transfected with 10 ng of GFP mRNA LNP and LNP$^{HA}$ with or without bsAb$^{HA-Suntag}$. Analysis of GFP expression was performed 24 hours post-transfection. (n=3 per group, 3 independent experiments performed)

c. Histogram showing GFP MFI of PDL1+ cells treated with the indicated bsAb:LNP weight ratio. 10 ng of GFP mRNA LNP was used in this experiment. Analysis of GFP expression was performed 24 hours post-transfection. (n=3 per group, 3 independent experiments performed)

d. DLS data of LNP$^{HA}$ before and after incubation with different ratio of bsAb:LNP$^{HA}$ as in **e.**

e. DLS data of LNP$^{HA}$ with different amount of total DSPE-PEG-HA. Histograms showing percentage of GFP positive K562$^{PD-L1/Suntag}$ cells in which GFP LNP$^{HA}$ and bsAb$^{HA-Suntag}$ were added at the same time or pre-incubated for 15 minutes prior treatment. Analysis of GFP expression was performed 24 hours post-transfection.

f. Histogram showing GFP MFI of PDL1 positive and negative cells treated with 20 ng of GFP mRNA LNP and LNP$^{HA}$ with different amount of DSPE-PEG-HA. Analysis of GFP expression was performed 24 hours post-transfection. (n=3 per group, 3 independent experiments performed)

**Supplementary Figure 3.**

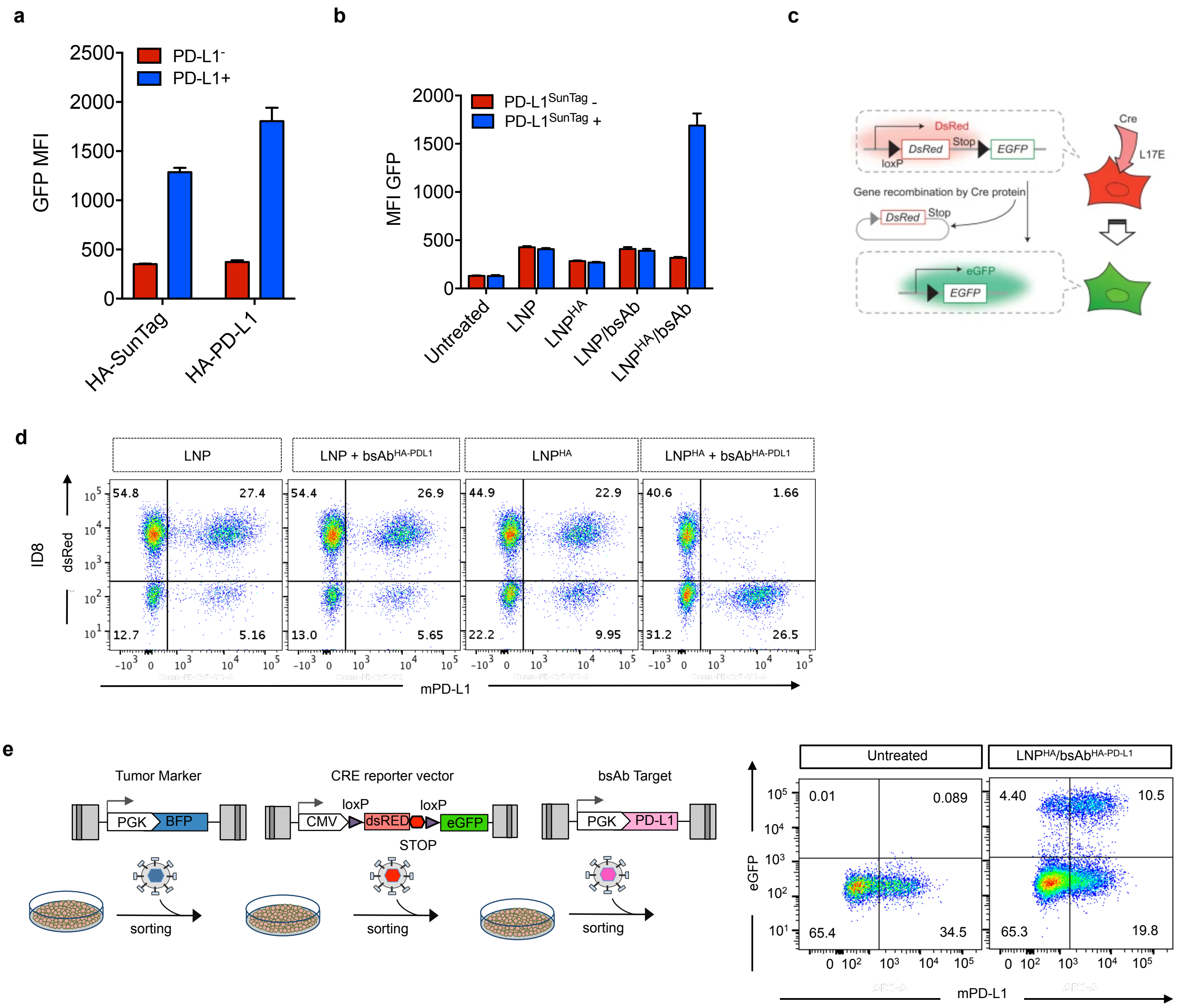

**Supplementary Figure 3**.

a. Histogram showing the MFI of GFP expression of $K562^{PDL1\text{-}Suntag}$ treated with eGFP $LNP^{HA}$ with $bsAb^{HA\text{-}Suntag}$ and $bsAb^{HA\text{-}PDL1}$. Cells were analyzed 24 hours post treatment. (n=3 per group, 2 independent experiments performed)

b. Histogram showing the MFI of GFP expression of $K562^{mPDL1}$ treated with eGFP LNP and $LNP^{HA}$ with or without $bsAb^{HA\text{-}PDL1}$. Cells were analyzed 24 hours post treatment. (n=3 per group, 2 independent experiments performed)

c. Schematic of the DsRed-STOP-loxP-eGFP cre recombination system.

d. Representative dot plot of ID8 cells engineered with DsRed-STOP-loxP-eGFP cre recombination system and treated with cre mRNA LNP and $LNP^{HA}$ with or without $bsAb^{HA\text{-}PDL1}$. Cre recombination analysis was performed 72 hours post-treatment. (n=3 per group, 1 independent experiments performed)

e. Schematic showing the lentiviral transduction strategy to generate tumor reporter cell lines for in vivo studies. B661, ID8 and B16F10 murine cell line were transduced with a BFP marker, $GFP^{stop}$ LoxP reporter system and mPD-L1. *Right:* Representative dot plots of $B16F10^{mPD\text{-}L1}$ engineered as the schematic, untreated and transfected with SM102-$LNP^{HA}$/$bsAb^{HA\text{-}PD\text{-}L1}$ complexes

### Supplementary Figure 4.

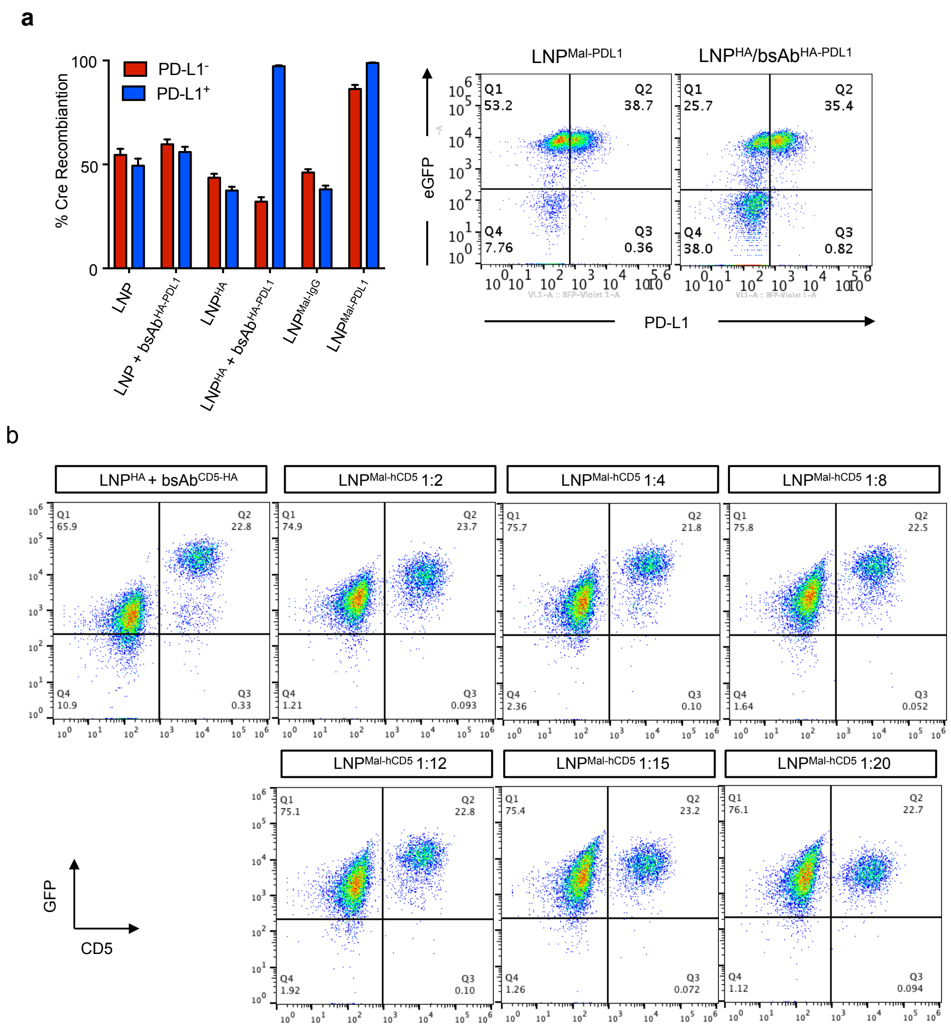

**Supplementary Figure 4**.

a. Left:Histograms showing the percentage of Cre recombination of K562$^{mPD-L1}$ treated with Cre LNP and LNP$^{HA}$ with or without bsAb$^{HA/PD-L1}$ and with LNP-IgG and LNP-PDL1. GFP expression was measured 72 hours post LNP transfection. Right: Representative dot-plots of K562$^{mPD-L1}$ treated with LNP-PDL1 and LNP$^{HA}$/bsAb$^{HA-PDL1}$. Note how treatment with LNP$^{HA}$/bsAb$^{HA-PDL1}$ is more specific than treatment with LNP-PDL1. Right: Representative dot-plots of K562$^{LoxP_GFP_mPDL1}$ treated with 1 ng of cre SM-102 LNP and LNP$^{HA}$ with or without bsAb$^{HA/PD-L1}$ and with bsAb$^{HA/ratIgG2b}$ with or without PD-L1 Ab. GFP expression was measured 72 hours post LNP transfection. (n=3 per group, 3 independent experiments performed)

b. Representative dot-plots of K562$^{CD5}$ cell line treated with LNP$^{HA}$ and bsAb or Mal-hCD5 conjugated LNP with different moral ratio of ab. Analysis was performed 24 hours post transfection. (n=3 per group, 3 independent experiments performed)

#### Supplementary Figure 5.

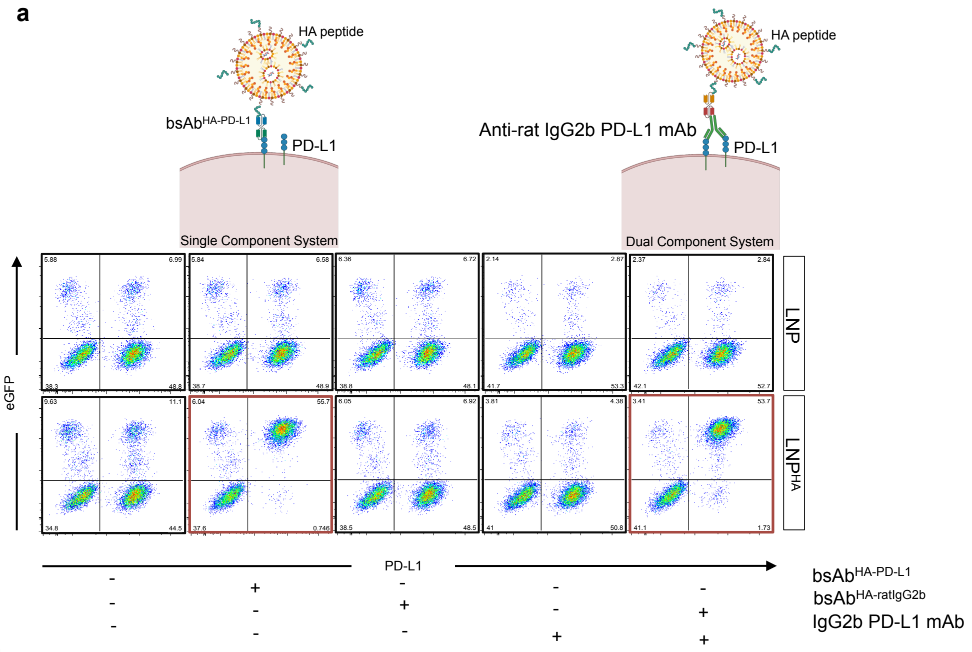


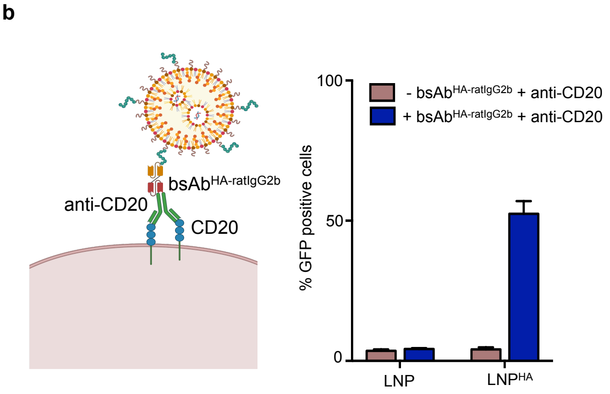

**Supplementary Figure 5**.

a. Representative dot-plots of $K562^{LoxP\_GFP\_mPDL1}$ treated with 1 ng of cre SM-102 LNP and $LNP^{HA}$ with or without $bsAb^{HA/PD\text{-}L1}$ and with $bsAb^{HA/ratIgG2b}$ with or without PD-L1 Ab. GFP expression was measured 72 hours post LNP transfection. (n=3 per group, 3 independent experiments performed)

b. Left: Schematic of $LNP^{HA}$/$bsAb^{HA\text{-}ratIgG2b}$/anti-CD20 Ab complex binding to CD20. Right: Histogram showing the percentage of eGFP positive BCL1 cells treated with 50 ng of eGFP SM-102 LNP and $LNP^{HA}$ with $bsAb^{HA/ratIgG2b}$/anti-CD20 Ab complexes. eGFP expression was measured 24 hours post transfection. (n=3 per group, 3 independent experiments performed)
